# Conformational flexibility, lipid binding, and regulatory domains in CelTOS

**DOI:** 10.1101/2022.03.31.486524

**Authors:** Hirdesh Kumar, John R. Jimah, Francis B. Ntumngia, Santosh A. Misal, Samantha J. Barnes, Michal Fried, John H. Adams, Paul H. Schlesinger, Niraj H. Tolia

## Abstract

CelTOS is an essential *Plasmodium* traversal protein and conserved in apicomplexan parasites. We showed that CelTOS forms pores in cell membranes to enable traversal of parasites through cells (Jimah et al., 2016). Here, we establish roles for the distinct regions of CelTOS, examine the mechanism of pore formation and evaluate the immunogenicity of engineered CelTOS variants. CelTOS dimer dissociation is required for pore formation as disulfide bridging between monomers inhibits pore formation and this inhibition is rescued by disulfide-bridge reduction. A helix destabilizing Pro127 allows CelTOS to undergo significant conformational changes to assemble into pores. The flexible C-terminus of CelTOS is a negative regulator that limits pore formation. Lipid binding is a pre-requisite for pore assembly as mutation of a phospholipid binding site in CelTOS resulted in loss of lipid binding and abrogated pore formation. The disulfide-locked and N-terminal deletion mutants showed improved immunogenicity relative to wild-type CelTOS in mice. These findings have implications for pore-forming proteins that are essential for diverse functions, identify critical regions in CelTOS, and will guide the design of effective CelTOS-targeting vaccines to combat infection and transmission of malaria and apicomplexan parasites.

## Introduction

Apicomplexan parasites infect a wide range of hosts and affect millions of humans and agricultural animals (Levine et al., 1980). Apicomplexan parasites include: *Plasmodia spp.,* the most severe parasite in the category that causes malaria (Black et al., 2010; Price et al., 2007); *Toxoplasma,* an opportunistic parasite that can cause severe toxoplasmosis in immunocompromised individuals; *Babesia,* the causative agent of babesiosis (Homer, Aguilar-Delfin, Telford, Krause, & Persing, 2000); *Theileria* (Shaw, 2003), a tick-born parasite that causes theileriosis in cattle; and *Cytauxzoon felis,* which causes cytauxzoonosis, an emerging tick-born disease in cats (Sherrill & Cohn, 2015). Among these apicomplexa-caused diseases, malaria results in the most significant mortality and morbidity, causing over 600,000 deaths and 200 million cases worldwide annually (WHO, 2019).

Apicomplexan parasites exhibit diverse biology and infect diverse hosts but share common features such as the presence of a definitive cell structure (the apical complex), a nonphotosynthetic plastid (the apicoplast), and actin-based motility. Cell traversal is another important common feature shared by most but not all apicomplexan parasites. Parasites traverse several cell types to complete essential segments of their life cycle inside the host and the vector. <u>Cell</u>-<u>t</u>raversal protein for <u>o</u>okinetes and <u>s</u>porozoites (CelTOS), Sporozoite Protein Essential for Cell Traversal 1 (SPECT1), and SPECT2 have established roles in cell traversal (Kumar & Tolia, 2019). Among these, CelTOS is relatively unique because it is required for cell traversal in both the human host and the arthropod vector. In addition, CelTOS is highly conserved among arthropod-borne apicomplexan parasites (Jimah et al., 2016).

Disruption of the CelTOS gene in *Plasmodium berghei* causes a 200-fold reduction in infectivity during mosquito stage development, and the resultant salivary gland sporozoites show defects in their ability to migrate through different cell types (Kariu, Ishino, Yano, Chinzei, & Yuda, 2006). Furthermore, CelTOS is a promising malaria antigen (Doolan et al., 2003) and immunization with recombinant CelTOS shows humoral and cellular responses against the parasite (Bergmann-Leitner et al., 2013; Bergmann-Leitner, Legler, Savranskaya, Ockenhouse, & Angov, 2011; Bergmann-Leitner et al., 2010). Finally, Amazonian populations elicit natural immune responses against *P. vivax* CelTOS (Rodrigues-da-Silva et al., 2017). These findings clearly highlight the importance of CelTOS in malaria parasites and in the human host response and confirm the potential of CelTOS as a malaria vaccine candidate.

Previously, we demonstrated that the structure of *P. vivax* CelTOS revealed that the helical organization in CelTOS resembled other membrane-disrupting proteins (Jimah et al., 2016). Soluble CelTOS exists as a dimer in which the two monomers protect the hydrophobic core and expose their hydrophilic residues at the surface (Jimah et al., 2016). CelTOS specifically binds to phosphatidic acid (PA) and forms pores in the inner leaflet of the host membrane (Jimah et al., 2016). However, how the CelTOS dimer rearranges and incorporates into the cell membrane is not clearly understood. Lack of information on the functional regions of CelTOS hinders structure-based design of effective CelTOS immunogens.

In this study, we report how the CelTOS dimer in solution transitions into a CelTOS pore in membranes and identify primary, secondary, tertiary and quaternary changes in CelTOS structure that regulate its fundamental function in pore formation. We tested the hypothesis that the soluble CelTOS dimer is in an inactive state and undergoes conformational rearrangements to penetrate cell membranes and form pores. We generated distinct CelTOS mutants and studied their dynamics and functional characteristics using 500 ns accelerated molecular dynamics (aMD) simulations and pore-forming assays. We developed a dimer-locked CelTOS using structure-based engineering to introduce cysteines at specific locations in CelTOS that prevent dimer dissociation. The disulfide-stabilized CelTOS dimer was inactive in the pore-forming assay. We identified a kink in helix 4 of CelTOS caused by Pro127, a residue conserved in other apicomplexans. Mutation of Pro127 to alanine stabilizes CelTOS and reduces its pore-forming activity, suggesting Pro127 may provide inherent instability to the CelTOS dimer. We further truncated the flexible N- and C-terminal segments individually, and observed an enhanced pore-forming activity upon deleting the C-terminal segment. On the contrary, deletion of the N-terminal has no significant difference from the wild-type protein. Finally, we showed Lys122 and Leu164 that comprise two conserved lipid-binding residues that interact with a likely phospholipid were essential for pore-forming activity. These data suggest the soluble CelTOS dimer undergoes a functionally essential conformation change upon contact with the lipid membrane. The disulfide-locked CelTOS and N-terminal truncated CelTOS elicited stronger antibody titers in immunized mice. These insights will inform the rational design of novel therapeutics targeting CelTOS against apicomplexan parasites.

## Results

### Engineered Cys residues stabilize CelTOS in an inactive, dimer state

We proposed that CelTOS is inactive in the dimer state observed in the crystal structure (PDB ID : 5TSZ) and undergoes significant conformational changes to form pores and incorporate into the inner leaflet of host cell membranes (Jimah et al., 2016). To test this hypothesis, we incorporated disulfide bonds in CelTOS that lock the dimer and prevent dimer dissociation. The CelTOS structure (PDB ID: 5TSZ) was analyzed for potential disulfide bridging residues, and Ser55 and Ala123 were identified; when mutated to cysteine (Ser55Cys and Ala123Cys), these residues formed a stable disulfide bridge (Figure 1A) (Craig & Dombkowski, 2013).

**Figure 1.**
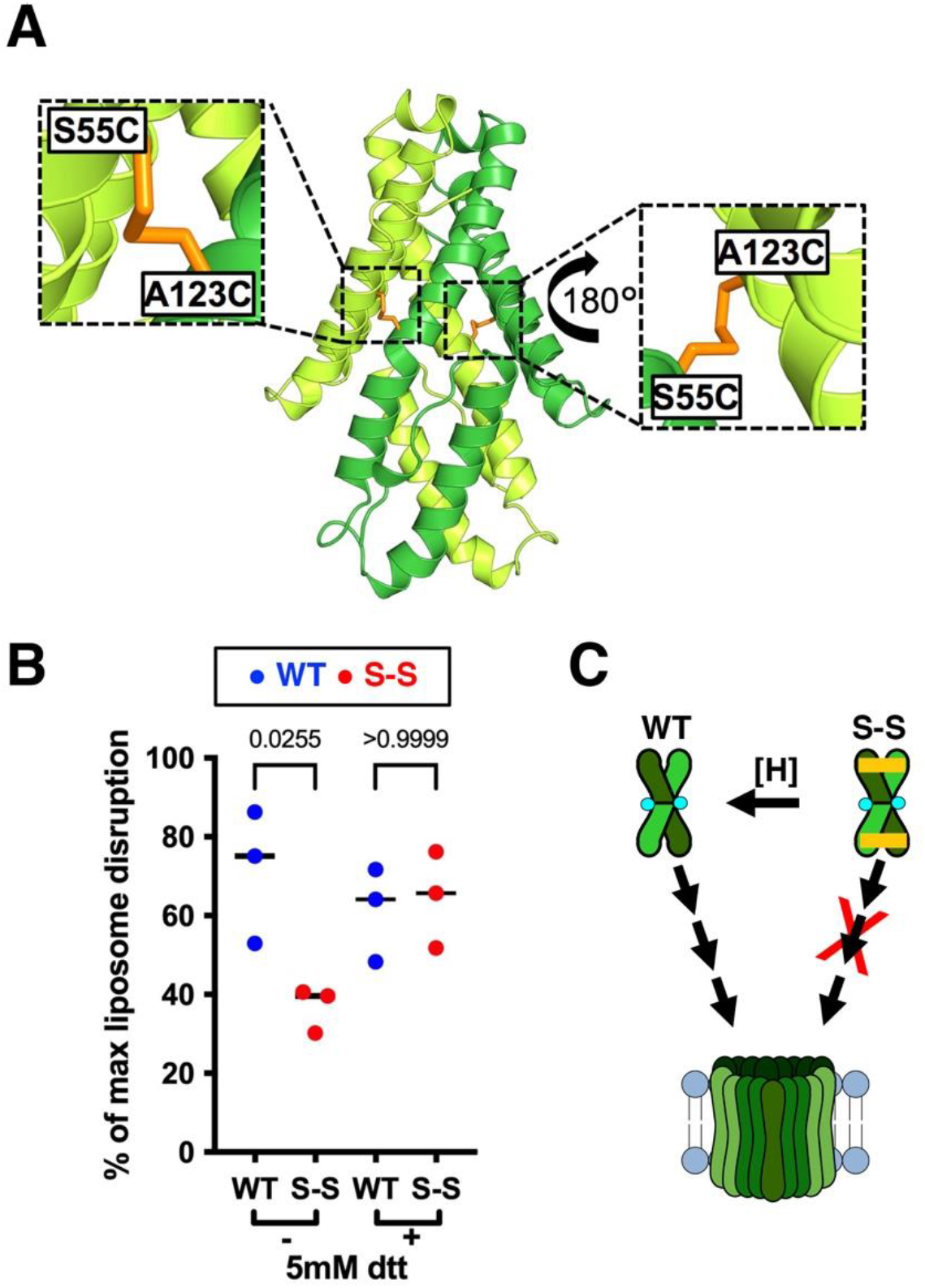
Disulfide-locked CelTOS dimer is inactive in pore formation. *(A)* CelTOS dimer (PDB ID: 5TSZ) is shown in the cartoon highlighting the location of Ala123 and Ser55. Note that the shown modeled structure highlights residues that were computationally mutated to cysteines to lock CelTOS in the dimer state. *(B)* Pore-forming assay: Disulfide-locked CelTOS dimer (S-S) is significantly less active than the wild-type protein (WT). Note that this loss of activity was rescued in presence of 5 mM dithiothreitol (DTT) that reduces the disulfide bridge, enables dimer dissociation and facilitates pore formation. The graph represents median values of three independent biological replicates each with 3-4 technical replicates (see Figure 1 – figure supplement 2 for three individual biological replicates demonstrating the reproducibility of the data). Significance was determined using Kruskal-Wallis analysis and Dunn’s multiple comparison. *(C)* A model depicting the plausible mechanism of conversion of soluble CelTOS dimer to the multimeric membrane pore (left) ; S-S inactivation (right) and its rescue by disulfide reduction (shown as [H]).

The disulfide mutant CelTOS (referred to as S-S) was readily expressed using a similar procedure as for wild-type CelTOS (WT) and the size exclusion chromatography (SEC) profiles of S-S and WT were indistinguishable (Figure 1 – figure supplement 1). A non-reduced SDS-PAGE analysis that retains disulfide bridging in the analysis but denatures the protein structure was conducted to evaluate the disulfide bridging of the various proteins (Figure 1 – figure supplement 1). This analysis revealed a band of ~37 kDa comprising a disulfide-locked dimer for the S-S mutant and a band of ~18 kDa corresponding to the uncrosslinked protein for WT. Together, the size-exclusion profile and the SDS-PAGE analysis suggest that the incorporated disulfide linkages create a cross-linked dimer but do not affect the native oligomeric state or structure of CelTOS dimer.

One concern of the introduction of the disulfide bridges is that CelTOS may form the expected disulfide bridges (C55-C123) of the parallel dimer but could also form the alternate disulfide bridges (C55-C55 or C123-C123) of an anti-parallel dimer. We investigated which disulfide bridging patterns were observed using mass spectrometry-based bottom-up proteomic analysis. In non-reduced conditions, only the interchain disulfide linkage between Cys55 and Cys123 (SGSTASSSLEGGSEF^55^CER-^123^CALEPTEK) was detected with a precursor abundance of 70 ± 7 % cross-linked species and 30 ± 7 % non-crosslinked species (Figure 1 A and Figure 1 – figure supplement 2). In three independent biological replicate experiments with independent purifications of the proteins, no disulfide linkage between Cys55-Cys55 or Cys123-Cys123 of the potential antiparallel dimer was detected, thus ruling out the possibility of alternate dimer organization in the S-S variant (Figure 1 – figure supplement 2). Together, with the SEC profile, the mass spectrometry analysis demonstrates the presence of the WT-like parallel dimer state in the S-S mutant.

We examined the pore-forming activity of the S-S mutant in non-reducing conditions and observed significantly less activity than the WT protein (Figure 1B and Figure 1 – figure supplement 3). Interestingly, this loss of pore-forming activity of the S-S mutant could be rescued in reducing conditions (5mM dithiothreitol) that eliminate the disulfide linkage (Figure1B and Figure 1 – figure supplement 3). These results indicate the addition of the cysteine residues has no effect on CelTOS function unless the cysteines are cross-linked to create a stabilized dimer.

Despite engineered disulfide linkages in the S-S mutant, the purified protein was a mixture of locked dimer (S-S) and unlocked dimer (WT) (Figure 1 – figure supplement 1) and therefore the S-S mutant showed residual membrane disruption activity. Indeed, mass spectrometry analysis of the non-reduced S-S dimer when the protein sample was alkylated with iodoacetamide to modify free sulfides revealed over 25 carbamidomethylated peptide spectral matches as free sulfides suggesting ~30% of all S-S dimer variant have a single disulfide bond or remained entirely unlocked (Figure 1 – figure supplement 2). Taken together, these results suggest that, for parasite cell traversal, the native CelTOS dimer must undergo dimer dissociation prior to significant conformational rearrangement to form pores in the lipid bilayer (Figure 1C).

### P127 causes a kink in helix 4 of CelTOS and is important for pore-forming activity

CelTOS is always observed as a dimer in solution (Jimah et al., 2016). However, we demonstrated that covalently stabilizing the dimer state by disulfide bridging inactivates CelTOS and activity can be restored only by eliminating this artificial disulfide bridge. We hypothesized that CelTOS contains inherently unstable region(s) that enable conformational rearrangements required for pore assembly and incorporation into the cell membrane. We found that the two monomers of the CelTOS dimer are not identical and deviate structurally from helix 4 to the C-terminus. The conformation of helix 4 differs between the two CelTOS monomers when the N-termini of the CelTOS monomers are aligned (Figure 2 – figure supplement 1). A deviation of 5.7 Å is observed between the Cα atoms of Thr135 of the two monomers (Figure 2 – figure supplement 1). The structural mobility is caused by Pro127 in helix 4 of CelTOS which causes a bend in the helix. Pro127 is also conserved among different CelTOS orthologs in other *Plasmodium* species (Figure 2A and Figure 2 – figure supplement 1) suggesting a critical role in its physiological function. Therefore, we suspected that Pro127 may provide internal flexibility to CelTOS structure.

**Figure 2.**
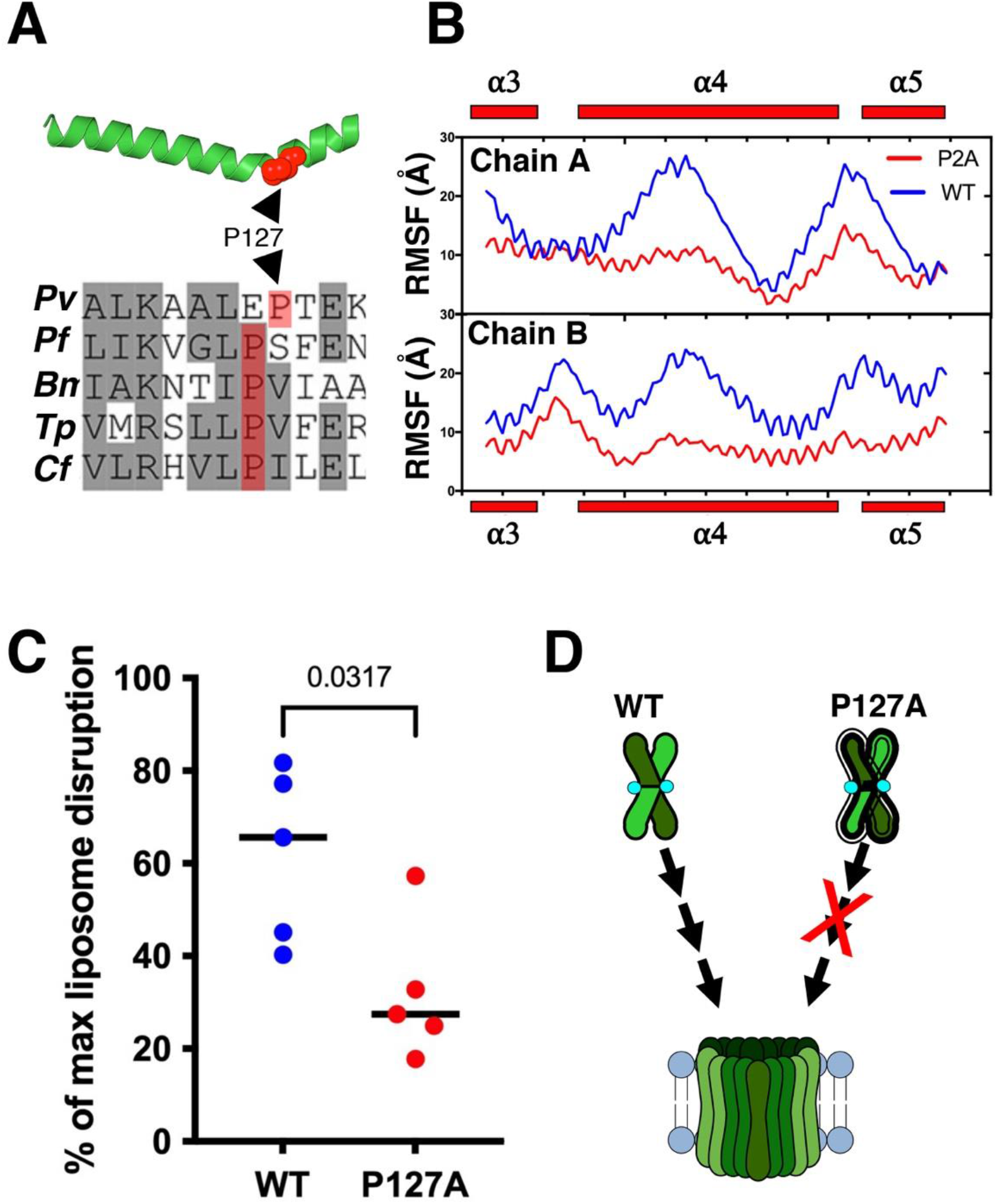
Pro127 is required for pore forming activity of CelTOS. *(A)* CelTOS helix4 (PDB ID: 5TSZ) is shown. Pro127 (shown in red sphere) causes a bend in the helix4 (top panel). Multiple sequence alignment (MSA) of CelTOS orthologs in apicomplexan parasites showing conserved Pro127 (highlighted in red). Residues in grey background represent conserved residues, and residues in white background represent non-conserved residues. *Pv*: *Plasmodium vivax, Pf: Plasmodium falciparum, Bm: Babesia microti, Tp: Theileria parva, Cf: Cytauxzoon felis (*bottom panel*). (B)* Root-mean-square-fluctuation (RMSF) plot of two monomers indicates that the Pro127Ala (P127A) mutant is less flexible in different regions including helix 4 (α4). *(C)* Pore-formation assay of CelTOS-wild type (WT) and Pro127Ala (P127A) mutant. The graph represents median values of five independent biological replicates each with eight technical replicates (see Figure 2 – figure supplement 2 for five independent biological replicates demonstrating the reproducibility of the data). Significance was determined using Mann-Whitney U-test. *(D)* A model showing the enhanced rigidity in P127A mutant causes loss of pore-forming activity.

We mutated Pro127 to Ala (P127A mutant), which was readily expressed with a SEC profile similar to the WT protein, suggesting P127A exhibits a similar architecture to WT (Figure 1 – figure supplement 1). In 500 ns aMD simulations, the P127A mutant was less flexible than the WT protein as evident from the lower root-mean-square-fluctuation (RMSF) values (Figure 2B). The P127A mutant showed a significant reduction in pore-forming activity compared to the WT protein, consistent with flexibility as a requirement for activity (Figure 2C and Figure 2 – figure supplement 3). The aMD simulation and pore-forming assay results imply that Pro127 provides flexibility to the CelTOS structure, which is functionally essential to provide conformational changes for CelTOS to incorporate into the inner leaflet of the host cell membrane (Figure 2D).

### Truncation of the flexible C-terminal residues increases CelTOS activity

The C-terminal disordered region lacks a transmembrane domain or sequence similarity to any known membrane-interacting domains (Figure 3A). When the two CelTOS monomers were aligned with respect to the N-terminal region, we observed a shift of 11.1 Å between Cα atoms of Glu143, suggesting that the C-terminal region is the most flexible region in the CelTOS structure (Figure 3 – figure supplement 1). The structural resemblance of C-terminal helices to membrane disrupting proteins (Jimah et al., 2016) and its flexibility in the CelTOS structure suggest the C-terminus may play a role in membrane penetration. We designed a C-terminal truncation mutant (C-del) comprising Leu36-Leu164 that omits residues C-terminal to helix 5 (His165-onward) (Figure 3A and Figure 2 – figure supplement 1). The purified C-del mutant eluted later than WT protein during SEC (Figure 1 – figure supplement 1), suggesting either an alternate conformation or dissociation of the dimer into monomers. Strikingly, C-del showed a ~50% increase in pore-forming activity compared to the WT protein (Figure 3B and Figure 3 – figure supplement 2). Taken together, these results establish that the flexible C-terminal region is a regulator that constrains the major conformation rearrangements in CelTOS required for the pore-forming activity (Figure 3C).

**Figure 3.**
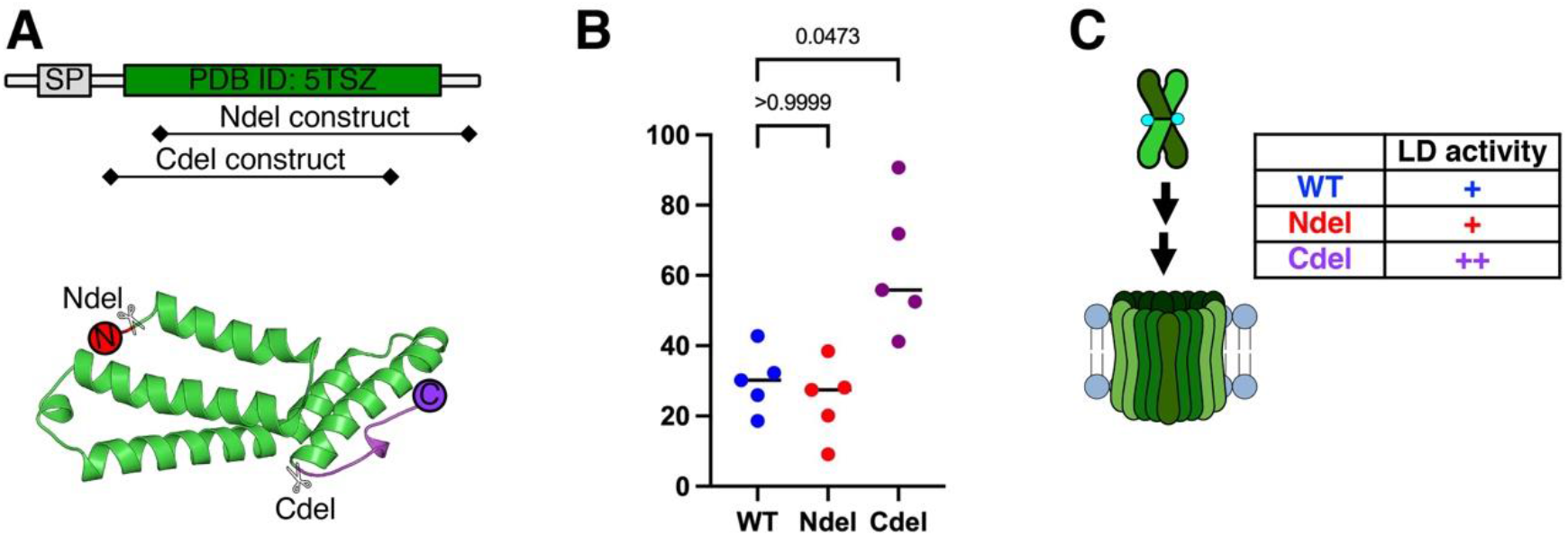
Truncation of flexible ends affect pore forming activity. *(A)* A schematic of *Plasmodium vivax* CelTOS protein sequence. Predicted domains are shown. SP=Signal peptide (*top*). Chain A of CelTOS dimer structure (PDB ID: 5TSZ) is extracted to show the regions of truncations highlighting flexible N- (red) and C- (purple) regions: N-del = Gly51-Asp196 ; C-del = Leu36-Leu164 *(bottom). (B)* Pore-forming activity of N-del and C-del mutants and their comparison to the WT protein. The graph represents median values of five independent biological replicates each with eight technical replicates (see Figure 3 – figure supplement 2 for five independent biological replicates demonstrating the reproducibility of the data). Significance was determined using Kruskal-Wallis analysis and Dunn’s multiple comparison. *(C)* A model comparing the pore-forming activity of the two mutants to the WT protein. While the N-del mutant (+) is similar to WT (+), the C- del mutant (++) shows enhanced pore forming activity.

To study the role of N-terminal residues, we designed an N-terminal truncation mutant that comprises residues Gly51-Asp196 (N-del) (Figure 2 – figure supplement 1) (Fig. 3A). N-del was readily expressed in *E. coli* and eluted as a dimer as confirmed by SEC profile (Figure 1 – figure supplement 1). Among five different biological replicates, a modest but non-significant reduction in activity of the N-del mutant compared to WT was observed. Collectively, the pore-forming activity of N-del was statistically non-significant from the WT protein (Figure 3B-C and Figure 3 – figure supplement 2).

### The two lipid-interacting residues Lys122 and Leu164 are required for pore formation

CelTOS specifically binds to phosphatidic acid (PA) and the CelTOS structure (PDB ID: 5TSZ) contains a buried surface area of 3003 Å^2^ that protects a hydrophobic core (Jimah et al., 2016). We observed two regions of unmodeled electron density in this hydrophobic core that could accommodate bidentate lipid molecules (Figure 4A). We hypothesized that these two unmodeled densities may correspond to two lipid molecules and therefore, the CelTOS-dimer may contain two lipid-binding sites.

**Figure 4.**
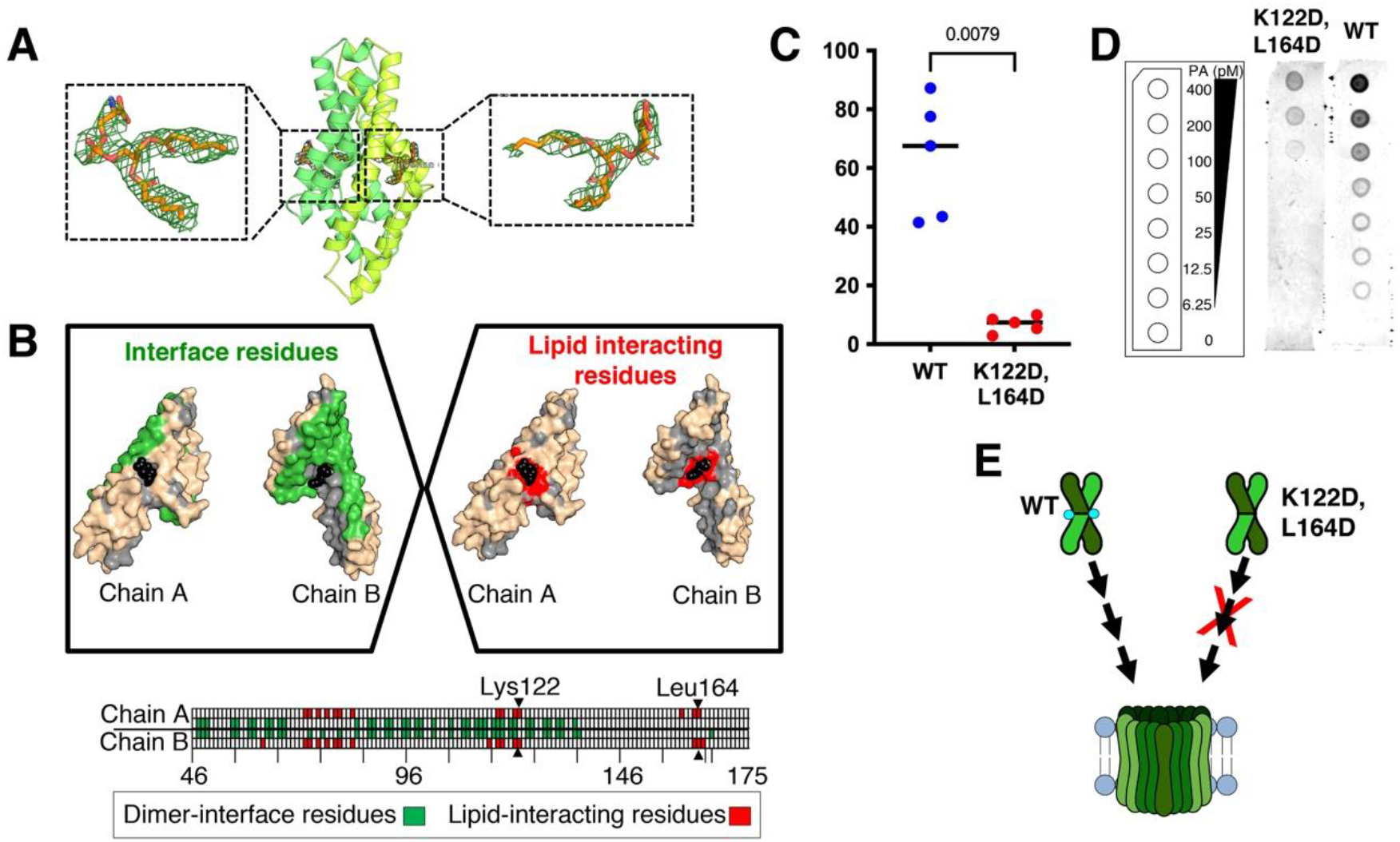
Lys122 and Leu164 are important for pore-forming activity of CelTOS. *(A)* Cartoon view of CelTOS structure (PDB ID: 5TSZ). The two monomers are shown in green and lime colors. Two POPA molecules (blue sticks) modeled in the electron densities are shown in the enlarged views. 2fo-fc map is contoured at 1.0 sigma. *(B)* Top: The two monomer structures of CelTOS showing the interface residues in green (left). CelTOS monomers showing the plausible lipid interacting residues in red (right), Bottom: An illustration of the CelTOS dimer showing the residues that are involved at the monomer-monomer interactions (green in color). Residues within 5 Å radius of bound lipid molecules are highlighted in red. Lys122 and Leu164 are shown. *(C)* Pore-formation assay of Lys122Asp,Leu164Asp(K122D,L164D) mutant and its comparison to the wild-type (WT) protein. The graph represents median values of five independent biological replicates each with eight technical replicates (see Figure 4 – figure supplement 2 for five independent biological replicates demonstrating the reproducibility of the data). Significance was determined using Mann-Whitney test. (*D*) Evaluation of binding to phosphatidic acid of WT and K122D,L164D mutant. Left: a schematic layout of the phosphatidic acid (PA) lipid strips containing a concentration gradient of PA from 400-0 pmol. Right: The double mutant shows poor binding to phosphatidic acid (PA). One representative biological replicate of three is shown. Two additional biological replicates are shown in Figure 4 – figure supplement 2. *(E)* A model showing the loss of pore-forming activity of the double mutant (K122D,L164D).

Two 1-palmitoyl-2-oleoyl-sn-glycero-3-phosphate (POPA) molecules were readily modeled into the electron density (Figure 4 A-B). Residues within a radius of 5 Å around the bound lipid molecules were then examined in detail. Among these, Glu73, Asn77 and Lys122 showed favorable polar interactions with the hydrophilic head group of POPA. Leu72, Ile75, Leu79, Ala80, Ile83, Leu117, Val118 and Leu121 showed non-polar interactions with the hydrophobic tails. These residues are conserved in CelTOS orthologs in different *Plasmodium* species (Figure 2 – figure supplement 1), suggesting CelTOS orthologs likely contain similar lipid-binding regions.

Most of these lipid-interacting residues are buried in the CelTOS dimer interface, and we suspected that attempts to manipulate these residues may affect the integrity and stability of the CelTOS dimer (Figure 4B). Therefore, we limited our approach to mutating lipid-binding residues that were located outside the CelTOS dimer interface. We identified Lys122, which makes hydrophilic interactions with the polar head of POPA and Leu164, which makes hydrophobic interactions with the non-polar tail of POPA.

The double mutant, Lys122Asp, Leu164Asp (referred to as K122D,L164D) was readily expressed in *E. coli.* The SEC profile of this mutant was similar to WT, suggesting the double mutations did not affect the dimeric state or the overall shape of the protein (Figure 1 – figure supplement 1). Next, we tested the pore-forming activity of K122D,L164D and found this double mutant was significantly less active than the WT protein (Figure 4C and Figure 4 – figure supplement 1). We have previously established a lipid-binding assay that revealed CelTOS can bind phosphatidic acid (Jimah et al., 2016; Jimah, Schlesinger, & Tolia, 2017). We leveraged this lipid binding assay to evaluate if the mutations in the lipid-binding region affect binding to phosphatidic acid. The K122D, L164D mutations showed a significant loss of lipid binding in comparison to the WT CelTOS (Figure 4D and Figure 4 – figure supplement 2). These results suggest Lys122 and Leu164 are functionally conserved lipid-binding residues essential for pore-forming activity of CelTOS (Figure 4E).

### S-S and N-del elicit stronger antibody titers in immunized BALB/c mice

Disulfide-linkage (S-S) or truncations (N-del, C-del) can significantly affect rearrangement and thus immunogenicity of the CelTOS protein. Therefore, immune responses against S-S, N-del and C-del variants were evaluated in BALB/c by injecting either WT CelTOS or engineered variants. A high-titer value of the CelTOS WT and other variants confirmed the functional efficacy of different injected antigens. When tested for induction of cross-reactive antibodies to wild-type, mice immunized with the S-S and N-del variants induced significantly higher antibody titers relative to the wild-type CelTOS. The C-del variant was the least immunogenic of all four antigens tested (Figure 5). These data indicate that structure-based design of CelTOS variants can improve the immune response upon vaccination.

**Fig 5.**
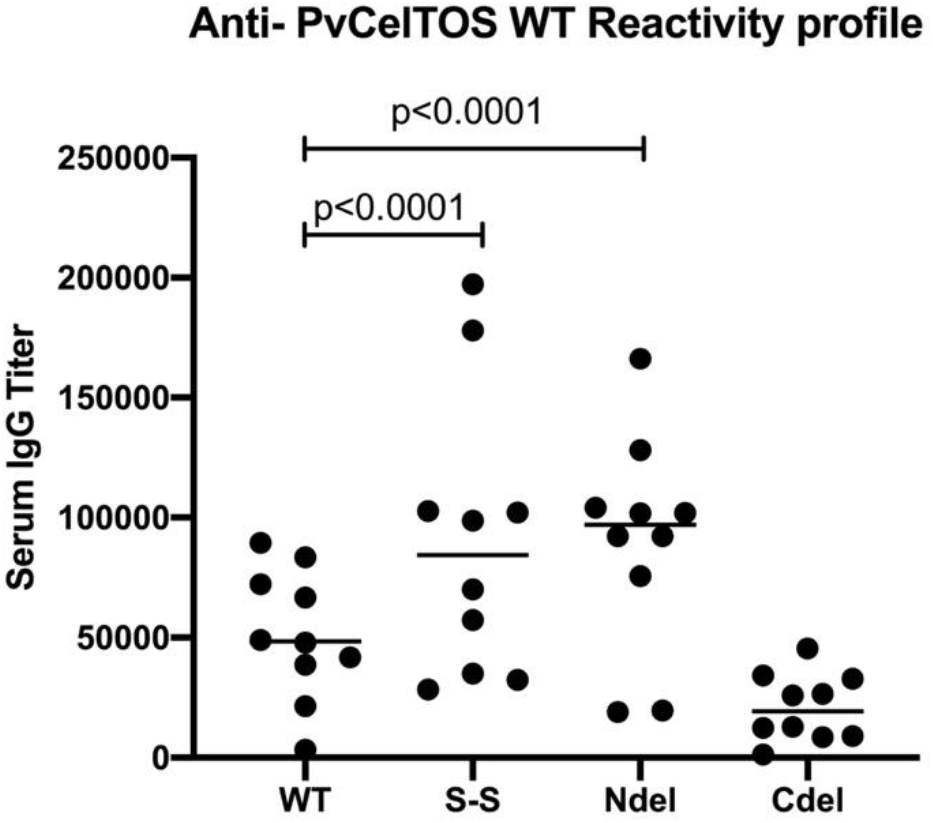
Anti-PvCelTOS (wild-type) reactivity profiles of different CelTOS variants. Immune sera raised against each recombinant PvCelTos antigen was evaluated by endpoint dilution ELISA for reactivity with wild-type antigen. Each data point represents IgG titers from individual mice serum (n=10), while lines represent median. The titers were determined as the serum dilution required to obtain an OD = 1.5. Kruskal-Wallis non-parametric ANOVA with a Dunnet’s Multiple Comparison with WT.

## Discussion

Malaria causes the highest burden of mortality and morbidity of all apicomplexan-caused diseases and affects millions of humans annually. *Plasmodium spp.,* the causative agent of malaria, undergoes a complex lifecycle within the mosquito vector and host. *Plasmodium* parasites must traverse several cell types within the vector and host before they find a suitable cell type to commit to development. Attesting to the crucial role of this cellular traversal, at least two classes of pore-forming proteins (PFP) are vital for *Plasmodium* growth and propagation (Guerra & Carruthers, 2017). Parasites lacking functional CelTOS expression are defective in mosquito and liver stage development (Kariu et al., 2006), suggesting CelTOS is essential and a promising therapeutic or vaccine target that can both block transmission to mosquitoes and prevent infections in humans.

PFP exist in a vast diversity of organisms but consistently require interaction with a target cell membrane, often in a receptor-specific manner, and undergo significant conformational rearrangement to penetrate into the membrane (Dal Peraro & van der Goot, 2016). This conformational rearrangement is required for the PFP to transform from a soluble to membrane-embedded protein (Anderluh & Lakey, 2008). In at least one case, the PFP is maintained in an inactive state by a bound chaperone, and membrane activation occurs upon chaperone release (Fraser, Karimi, Michalak, & Hudig, 2000). We have previously established that CelTOS specifically binds to phosphatidic acid and disrupts membranes through pore formation (Jimah et al., 2016). This suggests CelTOS needs neither a membrane bound receptor for its activity nor a chaperone to maintain its inactive state. Therefore, we hypothesized that the dimer state of CelTOS is an inactive state that must undergo significant conformational rearrangements to interact with and form pores in the lipid membrane. To test this hypothesis, we structurally designed a disulfide-stabilized dimer of CelTOS through incorporation of site-targeted cysteine substitutions. Valle *et al* used a similar disulfide bridge approach to lock sticholysin I in the homo-dimer state (Valle et al., 2018). The disulfide locked sticholysin I homo-dimer could not form pores and was 193 times less hemolytic than the WT protein (Valle et al., 2018). Similarly, Nguyen *et al.* created double-cysteine LukF (Leukocidin fast fraction) mutants and showed the disulfide-trapped mutants were inactive due to the transition arrest from pre-pore to pore (Nguyen, Higuchi, & Kamio, 2002). Similarly, the locked CelTOS dimer was inactive in pore formation. This loss of pore-forming activity was completely restored when the disulfide bond was reduced. These results clearly indicate that the soluble CelTOS dimer is indeed in an inactive conformation and establishes dimerization as a general method to regulate diverse pore-forming proteins.

Two monomers in the CelTOS dimer structure (PDB ID: 5TSZ) protect a hydrophobic core from the aqueous environment. We observed density in CelTOS structure that can represent two phosphatidic acid (PA)-containing lipid molecules in this position. The modeled PA molecules made polar and hydrophobic contacts with surface and core-region residues respectively. Such lipid-binding pockets have been reported to have a role in pore formation in other pore-forming proteins (K. Tanaka, J. M. Caaveiro, K. Morante, J. M. González-Mañas, & K. Tsumoto, 2015). For example, the monomeric, lipid-bound conformation of fragaceatoxin C (FraC), a pore-forming toxin protein, contains two conserved lipid-binding pockets which are primary sites of this molecule’s interaction with the cell membrane (Koji Tanaka, Jose MM Caaveiro, Koldo Morante, Juan Manuel González-Mañas, & Kouhei Tsumoto, 2015).

Purified CelTOS is always isolated as a dimer (Jimah et al., 2016), suggesting the dimer is the stable state in the aqueous solution. This raises immediate questions: why and how does this stable dimer undergo significant conformational rearrangement in contact with the lipid bilayer? To answer these questions, we analyzed the available CelTOS structure (PDB ID: 5TSZ) and identified a proline, Pro127, in the middle of helix 4. Prolines are well-known helix breakers(Cingolani, Petosa, Weis, & Muller, 1999) and, as expected, Pro127 causes a bend in the CelTOS helix 4. A similar proline-guided helix bend was been reported previously to confers the structural flexibility in importin-β(Kumeta, Konishi, Zhang, Sakagami, & Yoshimura, 2018). When these proline residues are mutated to alanine in importin-β, nuclear transportation is drastically reduced because the mutant’s structural flexibility is lost and cannot undergo rapid functional conformational changes (Kumeta et al., 2018). Membrane alignment of alpha-helices of toroidal pores demonstrate proline-induced bends play a critical functional role in stabilizing the pores (Schlesinger et al., 2002; Tuerkova et al., 2020). Similarly, in our current work, the CelTOS Pro127Ala mutant showed loss of pore-forming activity. The aMD simulation of Pro127Ala mutant showed less root-mean-square-fluctuation (RMSF) values compared to WT, suggesting a loss of structural flexibility in this mutant. The conservation of Pro127 in CelTOS orthologs in other *Plasmodium spp.,* as well as in highly divergent apicomplexan parasites, is strongly indicative that this residue has a functionally conserved mechanism for conferring inherent flexibility.

We examined the role of flexible N- and C-terminal regions in CelTOS. These flexible regions flank segments with structural similarity to membrane disrupting proteins (Jimah et al., 2016). To test our hypothesis if the N-terminal region assists CelTOS to undergo rearrangement, we truncated CelTOS before the start of helix 2 (between Gly51-Gly52). This mutant was readily expressed in bacteria but showed similar pore-forming activity to the WT protein. These results suggest the non-critical role of N-terminal residues in pore formation.

We analyzed further the potential functional importance of the C-terminal region of CelTOS by creating a C-terminal mutant protein by deleting His165-Asp196. Unexpectedly, this mutant showed a dramatic increase in pore-forming activity, suggesting the C-terminal region is a negative regulator to CelTOS activity. This finding is similar to our understanding of the functional importance of C-terminal regulatory domains defined in the BCl-2 family of PFPs (Cosentino & García-Sáez, 2017). The deletion of functionally homologous regulatory regions in a membrane fusion protein, <u>Snap</u> <u>re</u>ceptor (SNARE), promotes fusion of phospholipid endosomal vesicles (Parlati et al., 1999). Therefore, it is plausible that the malaria parasites may use the CelTOS C-terminal regulatory region to control the precise timing of pore formation and parasite egress from the host or vector cell (Figure 6).

**Figure 6.**
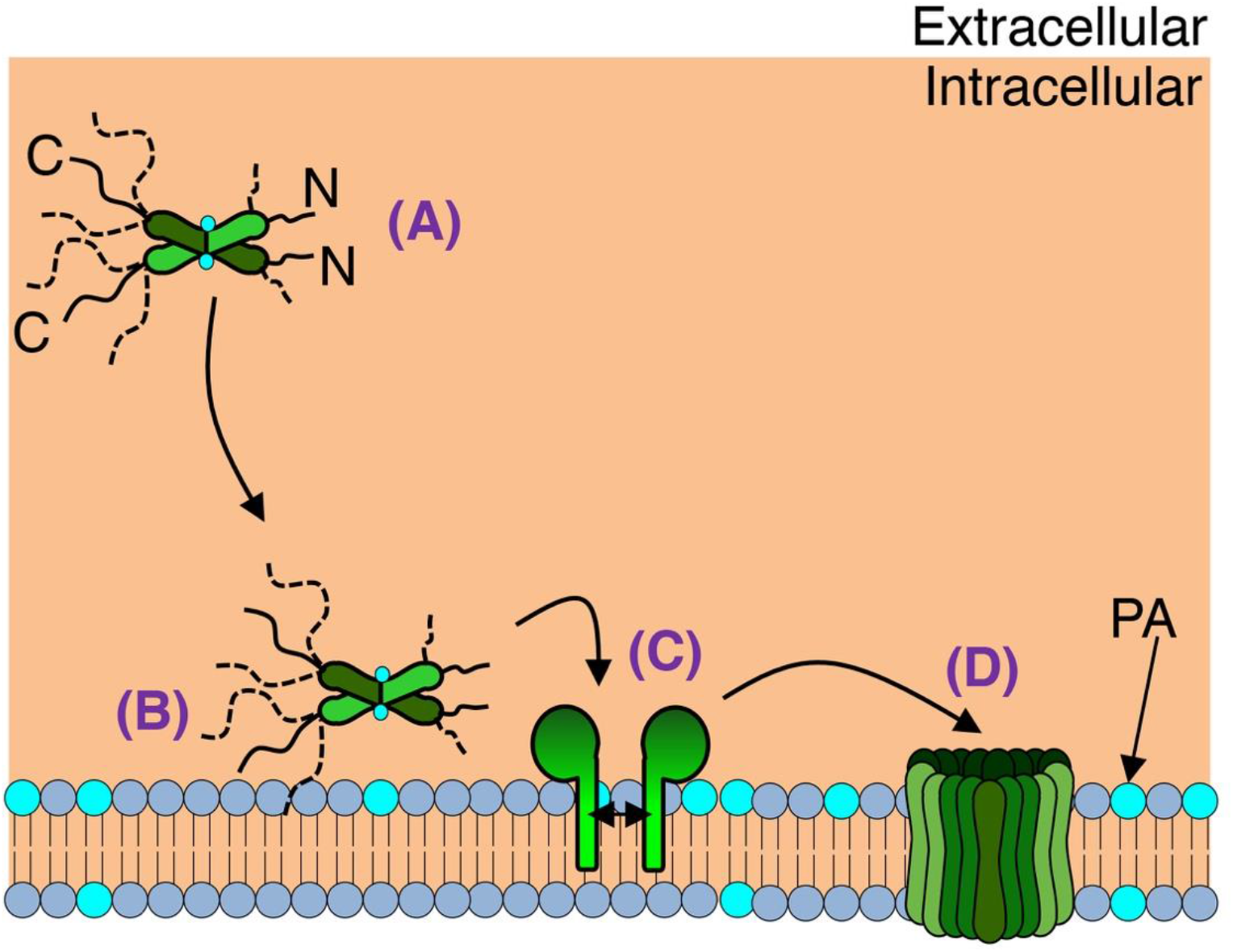
Proposed model for CelTOS-mediated pore formation. *(A)* CelTOS exists as a dimer in solution. Two lipid molecules can be observed bound to the CelTOS dimer. *(B)* Regulatory C-terminal domain (marked with “C” in panel A) prevents CelTOS from promptly interacting with the lipid bilayer. *(C)* Perpetual interaction between CelTOS and cell membrane leads to conformational rearrangement in CelTOS and thus penetration into the membrane. Membrane-inserted dimer dissociates into two monomers and forms pre-pore in the cell membrane. *(D)* Pre-pore leads to generation of mature pore.

CelTOS is localized on the *Plasmodium* surface and has an intracellular function in the host cell. CelTOS forms pores at the inner leaflet of the host cell and disrupts the host cell membrane (Jimah et al., 2016). Intracellular malaria parasites utilize CelTOS pores to exit the host cell. Plausible mechanisms of action of CelTOS-specific antibodies are either binding to the extracellular region of mature CelTOS pores (thus blocking the CelTOS-pore), or hitchhiking on the surface of an invading parasite to block surface-localized CelTOS (thus arresting parasite exit from the host cell).

In addition to studying the effect of different mutations on pore-forming activity of CelTOS, we felt encouraged to evaluate the immunization potential of different CelTOS variants compared to WT CelTOS. In particular, it became interesting to study the effect of different truncations (N-del, C-del) and conformational arrest (S-S) on antigen immunogenicity. We immunized BALB/c mice and measured antibody titers of immune sera raised against different CelTOS variants as well as WT CelTOS. Among three tested variants, immunized sera against the S-S variant showed the highest antibody titer against WT antigen, suggesting that conformational rigidity indeed enhances CelTOS-antibody interactions. Similarly, deletion of the flexible N-terminal residues plausibly enhances the conformational rigidity in the CelTOS-core domain, supported by our finding that the N-del variant induced improved antibody responses than WT CelTOS. Interestingly, the C-del variant was less immunogenic compared to WT CelTOS, suggesting that truncation of the flexible C-terminal region may result in a major conformational rearrangement in the CelTOS-core domain, thus diminishing antibody response against this variant.

Vaccine efficacy is typically better when higher titers are achieved. Therefore, immunogenicity of the different CelTOS variants, using the same mutations described above to study pore-forming activity of CelTOS, was evaluated relative to WT CelTOS. The significant differences in antigen immunogenicity (Fig. 5) suggests that conformational rigidity indeed enhances CelTOS-antibody interactions. Considering the highly divergent conformations WT-CelTOS can exhibit, structural design of the immunogen will be especially important going forward for improving the immunogenicity and functional efficacy of a CelTOS vaccine.

As a vaccine target *Plasmodium* CelTOS is a micronemal protein translocated to the parasite surface during the invasion process and would be directly accessible to soluble antibodies. However, the essential function of CelTOS in the sporozoite is intracellular during host cell traversal in the pre-erythrocytic liver stage infection. Nonetheless, the essential function of CelTOS to form pores at the inner leaflet of the host cell membrane allowing parasite exit from the host cell the (Jimah et al., 2016), is a process that can be inhibited by soluble antibodies (Bergmann-Leitner et al., 2010; Espinosa et al., 2017). While it is surprising that antibodies can inhibit intracellularly, we have similarly demonstrated anti-sporozoite antibody-mediated functional inhibition of sporozoite-liver stage intracellular development for antibodies to the *P. vivax* and *P. falciparum* circumsporozoite proteins, which are also micronemal proteins translocated to the sporozoite surface (Roth et al., 2018). Presumably, antibodies are readily hitchhiking on the surface of invading sporozoites. Further studies are needed to determine if anti-CelTOS inhibition that arrests sporozoite exit from the host cell functions by either blocking the CelTOS-pore forming function directly or indirectly.

In conclusion, our findings clearly demonstrate that CelTOS must undergo conformational rearrangements, including dimer dissociation, to form pores. We have shown that CelTOS contains lipid-binding pockets and these lipid-binding residues are conserved across apicomplexan parasites. We identified Pro127 as a conserved, key residue that provides inherent flexibility to CelTOS, assisting the required conformational rearrangement. We have shown that the C-terminal tail is a negative regulatory region in CelTOS. Among different CelTOS mutants, disulfide-locked and N-del mutants elicit high antibody titers against the WT CelTOS. These studies have widespread implications for the mechanisms of regulation and activation of pore-forming proteins and will inform the design of future generations of CelTOS-based vaccines.

## Materials and Methods

Multiple sequence alignment of CelTOS sequences from diverse apicomplexan parasites confirm that CelTOS is a cell traversal protein conserved in diverse apicomplexan parasites. To compare different CelTOS sequences, the following CelTOS sequences were retrieved from the Uniprot database (https://www.uniprot.org): *Plasmodium vivax* CelTOS (Uniprot id: A5JZX5), *Plasmodium falciparum* CelTOS (Uniprot id: Q8I5P1), *Babesia microti* (Uniprot id: I7J9F8), *Theileria parva* CelTOS (Q4N982), *Cytauxzoon felis* (PiroplasmaDB id: CF003135). These CelTOS sequences were submitted to the online Clustal Omega server (https://www.ebi.ac.uk/Tools/msa/clustalo/) in the FASTA format to perform multiple sequence alignment. Clustal Omega uses seeded guide trees and hidden Markov model (HMM) profile-profile techniques to perform a global alignment of sequences. The input protein sequences were submitted with the following default parameters: dealign input sequences: no, mbed-like clustering guide-tree: yes, mbed-like clustering iteration: yes, number of combined iterations: 0, max guide tree iterations: −1, max hmm iterations: −1, order: input. The percentage identity matrices and globally aligned sequences were downloaded.

To color the alignment files based on the conserved residues, the multiple-sequence-alignment file was submitted to the boxshade server (https://embnet.vital-it.ch/software/BOX_form.html). Boxshade is an algorithm that converts the input alignment files into the user-desired colored formats. The following parameters were used: output-format: RTF_new, Font-size: 10, consensus-line: none, Fraction-of-sequences: 0.5. The output grey-colored files were downloaded from the server.

Next, the secondary structure information was added to these aligned CelTOS sequences using *P. vivax* CelTOS structure (PDB ID: 5TSZ, www.rcsb.org) (Jimah et al., 2016).

### Cloning, expression and purification of different CelTOS mutants

#### Structure-guided design

*Plasmodium vivax* CelTOS coordinates (PDB ID: 5TSZ) were retrieved from the RCSB-PDB database and analyzed in PyMol (https://pymol.org/2/). CelTOS dimer (chain A and B) was extracted from the ternary complex and analyzed different protein regions in this dimer structure. The following mutants were shortlisted based on our structure-guided designs: Pro127Ala (to remove the bend in the helix4 region); Lys122Asp and Leu164Asp (to confirm the interaction of Lys122 and Leu164 interaction with the bound lipid); Ser55Cys and Ala123Cys (to lock the dimer through disulfide linkage); N-terminal truncation (Gly51-Asp196; to study the effect of N-terminal residues); C-terminal truncation (Leu36-Leu164; to study the effect of C-terminal residues).

#### Cloning

Previously cloned pET28+ vector that contains codon-optimized PvCelTOS (Gene ID: PVX_123510) sequence was used (L36-D196)(Jimah et al., 2016). This vector contains the T7 promoter followed by PvCelTOS sequence and a 6x-His tag at the C-terminus of PvCelTOS. Inserts were cloned using SalI and XhoI restriction sites.

#### Expression and purification

The cloned mutants were transformed in BL21 (DE3) competent *E. coli* cells, grown in the lysogeny broth (LB) media containing kanamycin (10 μg/mL) at 37° C. The culture was induced with 1mM isopropyl thio-ß-d-galactoside (IPTG) at an O.D. of 0.6-0.8 for 3h at 37° C.

To purify the CelTOS mutants, cell pellets were resuspended in lysis buffer (50 mM Tris-HCl, pH7.4, 150mM KCl, 5mM Imidazole) and lysed by sonication at 70% power for 3 minutes (0.5 sec: ON; 0.5 sec: OFF). The lysate was centrifuged at 10,000 xg for 20 min and the His_6_-tagged-proteins were purified by Ni-NTA chromatography followed by size exclusion gel filtration using Superdex 200 10/300 GL column (GE Healthcare Life Sciences, Pittsburg, PA).

### Pore-forming assay

1-palmitoyl-2-oleoyl-sn-glycero-3-phosphocho-line (POPC) and 1-palmitoyl-2-oleoyl-sn-glycero-3-phosphate (POPA) lipids (Avanti Polar Lipids, Alabaster, AL) were dissolved in chloroform, dried under N2 gas (20-30 minutes). Dried lipids were hydrated in diethyl ether hydrated in 10 mM HEPES pH 7.4, 150 mM KCl. To study appearance fluorescence upon dequenching 20 mM carboxyfluorescein (in 10 mM HEPES pH 7.4, 150 mM KCl) was added to the hydrated lipid solution. Liposomes were prepared by serial cycles of vigorous mixing and ultrasonic bath treatment. Thus prepared, quenched and 6-Carboxyfluorescein-filled liposomes were extruded through Whatman® Nuclepore™ Track-Etched Membrane (pore size=200 nm) (Sigma Aldrich, Saint Louis, MO) as previously described(Jimah et al., 2017). Finally, 6-Carboxyfluorescein filled liposomes were separated from the unincorporated 6-Carboxyfluorescein by size exclusion chromatography using Sephadex G 25–300 (Sigma Aldrich, Saint Louis, MO).

To study the effect of different mutations on CelTOS structure, we added varying concentrations (25uM, 100uM and 250uM) of each mutant to the 250 nM liposome solution. The solution was incubated for 5 min at room temperature and the carboxyfluorescein release from liposome was observed using a Fluorescent plate reader at 512 nm upon excitation at 492 nm. Finally, Triton-X-100 was added to complete dequenching of liposome in each experiment. Three replicates of each experiments were performed for all the mutants. The percent liposome disruption (LD) was calculated as:

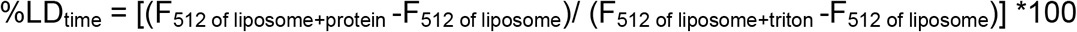

### Lipid blot assay

CelTOS binding to lipid strips were performed as previously described (Jimah et al., 2016; Jimah et al., 2017) and per manufacturer’s recommended protocol. Customized phosphatidic acid (PA) membrane strips (Echelon Biosciences Inc.) containing a concentration gradient of PA from 400-0 pmol (400 pmol, 200 pmol, 100 pmol, 50 pmol, 25 pmol, 12.5 pmol, 6.25 pmol, and 0 pmol) were incubated in blocking solution (50 mM Tris pH 8.0, 100 mM NaCl, 0.1% Tween 20, 3% BSA) for 1 hr at RT. Purified CelTOS variant was added to a membrane at a final conc. of 1 uM in blocking solution for 1 hr at RT. Membranes were next treated with 1: 1000 dilution of primary antibody (6-His tag mouse anti-tag, Invitrogen) for 1 hr at RT. The membrane was subjected to 1: 10 000 dilution of secondary antibody (IRDye® 800CW Goat anti-Mouse IgG Secondary Antibody; LI-COR Biosciences) in blocking solution for 1 hr at RT. The membrane was washed 3-times between each steps with wash solution (50 mM Tris pH 8.0, 100 mM NaCl, 0.1% Tween 20). Finally, the fluorescence signal was detected using Odyssey CLx imaging system.

### Molecular dynamics simulations

To study the effect of different mutants on CelTOS structure, we performed accelerated molecular dynamics simulation as described below.

#### In silico mutagenesis

Prior to mutagenesis, structure was prepared using Protein Preparation Wizard (Impact 6.3, Schrodinger 2014-2, Maestro 9.8)(Sastry, Adzhigirey, Day, Annabhimoju, & Sherman, 2013) as previously described (Sharma & Wakode, 2016). In brief, the structure was corrected for atoms and bonds, and energy minimized. Maestro visualizer was used to mutate the residues (in both the chains) and generate the following CelTOS mutants: Pro127Ala (Pro127 to Ala); Lys122Asp and Leu164Asp (K122D,L164D); S-S (Ser55/Ser123 to Cys); N-del (Leu36-Phe54 fragment was deleted); and C-del(His165-Asp196 fragment was deleted).

#### System preparations

All the designed mutants were individually prepared using tleap module of AMBER14 (Assisted Model Building with Energy Refinement). FF14SB force field parameters were set for the protein using the AMBER14 LEaP module (Hornak et al., 2006). The AM1-BCC method was used to assign partial atomic charges for bound inhibitor (POPA) and general amber force field (GAFF) was used to create its topology (Wang, Wolf, Caldwell, Kollman, & Case, 2004). Na^+^ ions were treated according to the “non-bonded” model method (Stote & Karplus, 1995). The prepared systems were solvated with TIP3P water model by creating an cubic water box, where distance of the box was set to 10 Å from periphery of protein (Kiss & Baranyai, 2011). Molecular systems were neutralized through the AMBER LEaP module by the addition of a necessary amount of counter ions (Na^+^) to construct the system in an electrostatically preferred position. Na^+^ and Cl^-^ ions were added to maintain the ionic strength of 150 mM. The whole assembly was saved as the prepared topology and coordinate files to use as input for the PMEMD module of the AMBER (Case et al., 2005).

#### System minimization, heating and equilibration

Prepared systems were energy-minimized in a two-step process: initial 1000 steps of steepest descent, during which each complex was fixed to allow water and ion movement, followed by 500 steps of conjugate gradient minimization of the whole system (complex, water and ions). The minimized systems were gradually heated from 0 to 298 K using an NVT ensemble for 100ps where the protein-ligand complex was restrained with a large force constant of 5 kcal/mol/Å^2^.

Following heating, the systems were equilibrated under constant pressure at 298 K and the restrain was gradually removed at NPT ensemble as follows: 5 kcal/mol/Å^2^ (40 ps), 2 kcal/mol/Å^2^ (20 ps), 1 kcal/mol/Å^2^ (20 ps) and 0.5 k cal/mol/Å^2^ (10 ps).

#### Initial conventional molecular dynamics (cMD)

Prior to an accelerated molecular dynamics simulation (aMD), a 50 ns long conventional molecular dynamics (cMD) simulation was performed to calculate the system-specific parameters. Each cMD simulation was performed on NPT ensemble at 298 K temperature and 1 atm pressure. The step size of 2 fs was kept for whole simulation study. Langev in thermostat and barostat were used for temperature and pressure coupling. The SHAKE algorithm was applied to constrain all bonds containing hydrogen atoms (Gunsteren & Berendsen, 2006). Non-bonded cutoff was kept on 10Å and long-range electrostatic interactions were treated by Particle Mesh Ewald method (PME) with fast Fourier transform grid spacing of approximately 0.1nm (Darden, York, & Pedersen, 1993).

#### Accelerated molecular dynamics (aMD)

Accelerated molecular dynamics is an all-atom enhanced sampling method to identify metastable conformational states in the protein structure. The following parameters were extracted from the initial cMD simulation to start a 500 ns aMD simulation for each system:

1. Average total potential energy threshold (EthreshP; kcal/mol) = Total potential energy (kcal/mol) + 0.16kcal/mol/atom x (number of total atoms)
2. Inverse strength boost factor for the total potential energy (alphaP) = 0.16 kcal/mol/atom x (number of total atoms)
3. Average dihedral energy threshold (Edih; kcal/mol) = 4 kcal/mol/residue x (number of solute residues)
4. Inverse strength boost factor for the dihedral energy (alphaD) = (1/5) x 4 kcal/mol/residue x (number of solute residues)

The last 200 ns production run was analyzed using the cpptraj module of the AMBER14 and VMD (Humphrey, Dalke, & Schulten, 1996). Trajectory snapshots were taken at each 100ps, which were used for final analysis. The minimization and equilibration were performed by PMEMD module of AMBER14. The production simulations were performed using PMEMD program of AMBER running on NVIDIA Tesla C2050 GPU work station (Gotz et al., 2012).

#### Analysis

The stability of each system during the simulation was studied by calculating the root-mean-square deviation (RMSD) of the backbone atoms of different frames to the initial conformation, and therefore RMSD is the measure of the average distance between the atoms (usually the backbone atoms) of superimposed protein structures. The stable trajectories were further processed for RMSF analysis.

Relative mean square fluctuation (RMSF)

To study the effect of different mutations on the overall protein flexibility, average fluctuation values were followed for individual amino acid residues during the simulations. The RMSFs were calculated with the backbone atoms of amino acid residues for each structure using the following formula:

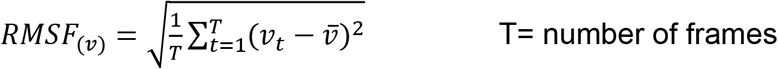

RMSF values were calculated using the CPPTRAJ module of AMBER 14.

### Abundance measurement of the disulfide-linked and non-linked peptides of S-S dimer by Mass spectrometry

#### Sample preparation and LC-MS/MS

Purified protein was buffer exchanged to 25mM ammonium bicarbonate (pH 7.8) and the protein concentration was measured by nanodrop at A280.

Approximately 10μg proteins were alkylated with 5mM iodoacetamide in dark for 30 min. Trypsin/Lys-C mix (Mass Spec Grade, Promega) was added to protein solution at 1:50 ratio (trypsin: protein) and incubated for 16hr at 37°C. The enzymatic digestion was terminated by adding 0.1% formic acid (FA). The tryptic digest was further purified by C18 Zip Tips (Millipore Sigma, USA) and reconstituted in 0.1% FA. The peptide solution was centrifuged at 15000 RPM before loading on a reversed-phase C18 column. Approximately, 800 ng of peptide solution was loaded onto the EASY-Spray PepMap RSLC C18 (15 cm × 150 μm) column in mobile phase A (0.1% FA in water (LCMS grade)) using UltiMate 3000 UHPLC System (Thermo Fisher Scientific, USA). The peptides were separated using a 60-min gradient from 2 to 30% mobile phase B (0.1% FA in Acetonitrile) from 5 to 40 min and ramped up the mobile phase B up to 90% for another 8 min and hold at 90% B for two minutes at a flow rate of 300 nL/min. The peptide eluent from the C18 column was electrosprayed in Thermo Orbitrap Fusion Lumos mass spectrometer (Thermo Fisher Scientific, USA) in positive ion mode using 2.2 kV voltage. Peptides with the precursor mass in the m/z range 375 to 1800 with 2 to 8 charges were selected for HCD activation (normalized collision energy 35%) with 60 S of dynamic exclusion. The resolution for MS1 was set at 120,000 (FWHM at m/z 200) and HCD fragment ions were analyzed with a resolution of 30,000 (FWHM at m/z 200). Samples were run in three independent biological triplicates.

#### LC-MS/MS data analysis

The raw data files were analyzed on Thermo Proteome Discoverer v2.5 (Thermo Fisher Scientific, USA) using SEQUEST HT and XlinkX/PD search nodes for the peptide and disulfide linkage search respectively. Enzyme digestion was set as full trypsin with a maximum of three missed cleavages. Mass tolerances for precursor and fragment ions were set to 5 ppm and 0.02 Da respectively. Variable modifications included methionine oxidation(+15.995 Da), and carbamidomethyl cysteine (+57.021 Da). A disulfide (−2.016 Da) linkage on cysteine residue was set as a crosslink modification in XlinkX/PD node. Peptides and crosslinks with a 1% false discovery rate (FDR) were selected for further analysis. Disulfide-linked peptides with a minimum XlinkX score of 40 were considered as confident identifications. The precursor abundance of peptides and disulfide-linked peptides were used for quantification. The sum of log_2_ transformed precursor abundance of each peptide and crosslinked species were plotted and the mean of three was used to calculate the percent abundance of disulfide-linked and non-linked cysteine residue containing peptides.

### Immunizations

This study was performed in strict accordance with the recommendations in the Guide for the Care and Use of Laboratory Animals of the National Research Council of the National Academies. All animals were handled according to approved institutional animal care and use committee (IACUC) protocols (#IS00004488 and #IS00008486), Division of Research Integrity and Compliance, University of South Florida.

Polyclonal antibodies were raised in female BALB/c mice (6-8 weeks old), obtained from Harlan Laboratories, Inc. and housed under specific pathogen-free conditions. Pre-immune sera were collected from each mouse prior to immunizations. Groups of mice (n=10) were immunized twice subcutaneously at three-week intervals with 25 μg/dose of either recombinant CelTOS (WT) or mutant alleles formulated in Titermax Gold adjuvant (TiterMax®), as previously reported (Ntumngia et al., 2013). A control cohort was immunized with PBS and adjuvant alone. Mice were exsanguinated three weeks after the second immunization and serum was separated and stored at −20°C until needed.

### Measurement of antibody titers to recombinant proteins

Mice sera were tested individually for reactivity with both homologous and heterologous recombinant CelTOS proteins by end point titration ELISA (Ntumngia et al., 2013). Briefly, 96 well micro titer plates (Maxisorb-Nunc) were coated over night at 4°C with 100 μL/well of recombinant CelTOS or mutant alleles, diluted to 2 μg/mL in coating solution (KPL). Unbound proteins were washed off with PBS/0.05% Tween 20 and any unbound surfaces blocked with 5% skimmed milk powder in wash buffer for 2 hr at room temperature. Then, plates were incubated for 2 hr at room temperature with 100 μL/well of 3-fold dilutions of mice sera (starting at 1:1000 dilution). The plates were washed, and wells incubated for 90 min at room temperature with 100 μL/well of AP-conjugated goat-anti-mouse (H+L) antibody (KPL Inc.) diluted 0.5 μg/mL in blocking solution and bound antibody detected with 100 μL of alkaline phosphatase substrate conjugate (KPL Inc.). The reaction was stopped after 20 min with 100 μL/well of APstop^TM^ solution (KPL Inc.) and absorbance read at 630 nm on a microplate reader (BioTek Instruments Inc). An anti-PvDBP sera was used as negative control, while pre-immune sera or sera from mice immunized with adjuvant alone was used as background control. Antibody titers were determined as reciprocal of serum dilution required to give an OD = 1.5 (Mullen et al., 2006).

### Statistical analyses and software

All analyses were performed using Prism 9 (GraphPad Software, La Jolla, CA).

## Acknowledgements

This work was supported by the Intramural Research Program and the Extramural Research Program (R01AI137162) of National Institute of Allergy and Infectious Diseases, National Institutes of Health and the Burroughs Wellcome Fund. The authors thank J. Patrick Gorres for editing the manuscript.

## Author Contributions

N.H.T., H.K. and J.R.J. conceived the study. N.H.T., H.K. and J.R.J. designed the experiments; H.K., J.R.J., S.A.M. and F.B.N. performed experiments; H.K., J.R.J., S.A.M., F.B.N. and S.B.J. analyzed data; J.H.A., M.F., P.H.S. and N.H.T. verified the analytical methods and supervised the findings; H.K. and N.H.T. wrote the manuscript with critical edits from P.H.S., M.F. and J.H.A. All authors provided critical feedback and helped shape the research, analysis and manuscript.

## Legends and figure supplement files

**Figure 1 – figure supplement 1:**
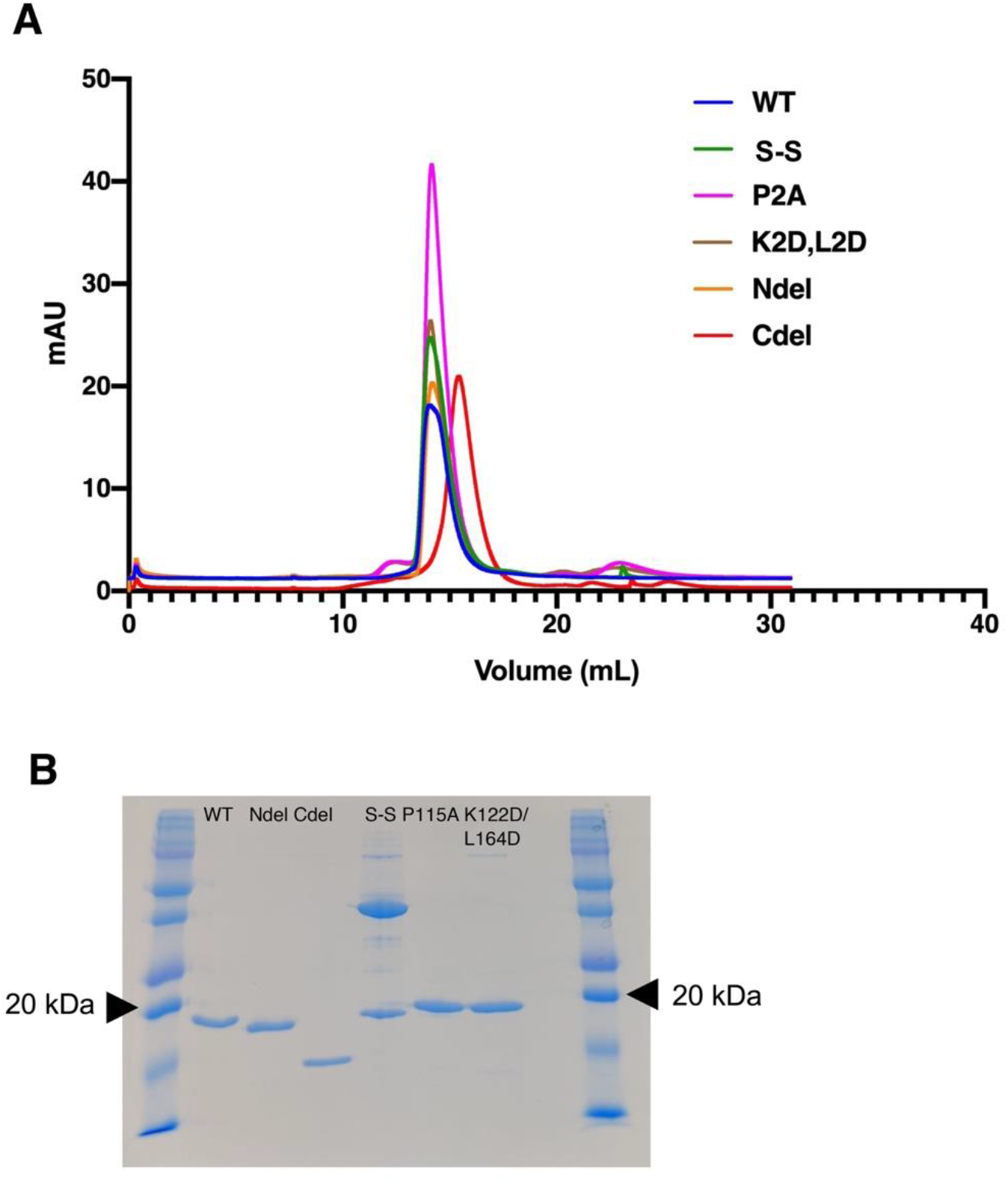
Comparison of size exclusion chromatography profiles of different CelTOS mutants. *(A)* Purified CelTOS WT (WT) or different mutants were independently loaded on size-exclusion column and eluted peaks are shown in different colors. *(B)* Peak fraction of each protein was run on 15% SDS-PAGE under non-reducing conditions that allow for disulfide bridges to be retained but protein structure to be denatured. The 20 kDa molecular weight marker is shown by arrow heads. All proteins and mutants are pure and show a single band at the expected molecular weight of the protein with the exception of the disulfide locked dimer that reveals a band at the expected molecular weight of a disulfide cross-linked dimer.

**Figure 1 – figure supplement 2:**
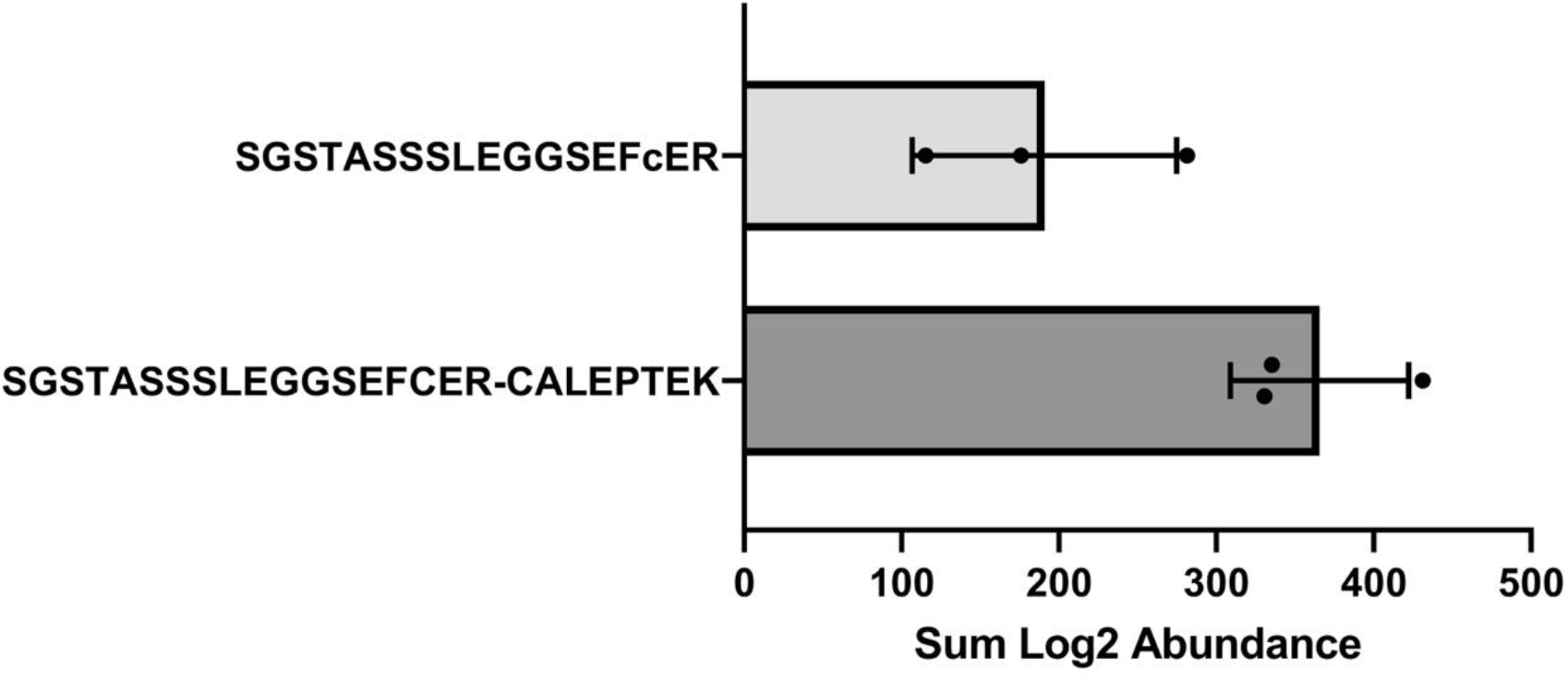
Precursor abundance measurement of the peptidic ions of disulfide-locked cysteines or free sulfides.

**Figure 1 – figure supplement 3:**
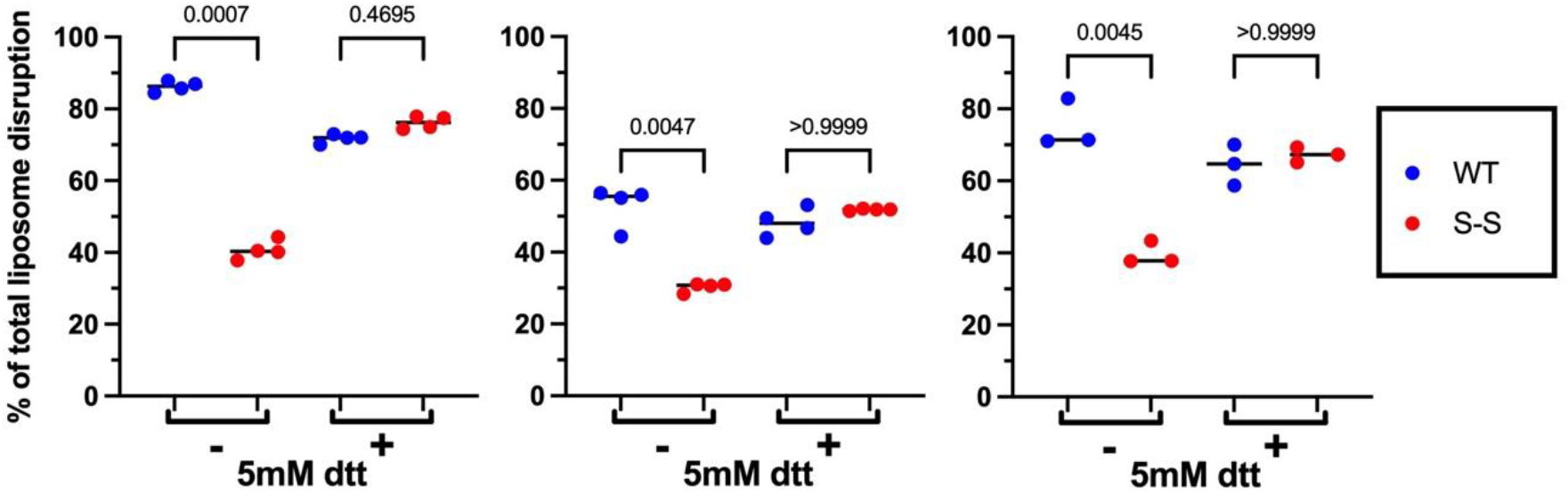
Pore-formation assay of disulfide-locked CelTOS dimer (S-S) and its comparison to the wild-type (WT) protein. S-S is significantly less active than the WT. Note that this loss of activity was rescued in presence of 5 mM dithiothreitol (DTT) that reduces the disulfide bridge and enables pore formation. Each graph represents an individual biological replicate that was performed on a separate day using freshly expressed and purified proteins and freshly prepared liposomes. Each biological replicate consists of three to four technical replicates. Lines represent median values. Significance within each biological replicate was determined using a Kruskal-Wallis analysis and Dunn’s multiple comparison. P-values are shown for each comparison. The y-axis is consistent between each graph and therefore is shown only for the left-most graph. The color-coding is same for each graph and therefore shown only for the right-most graph. The difference in absolute activity is likely due to inherent biological variability derived from each independent batch purification of protein or liposomes. The replicability of the data with independent preparations of proteins and liposomes and performed on different days demonstrate the pore-disruption assay is robust and can distinguish between WT and S-S proteins. The means of each biological replicate were used in Figure 1B to compare WT and S-S proteins across multiple days, batches and preparations.

**Figure 2 – figure supplement 1:**
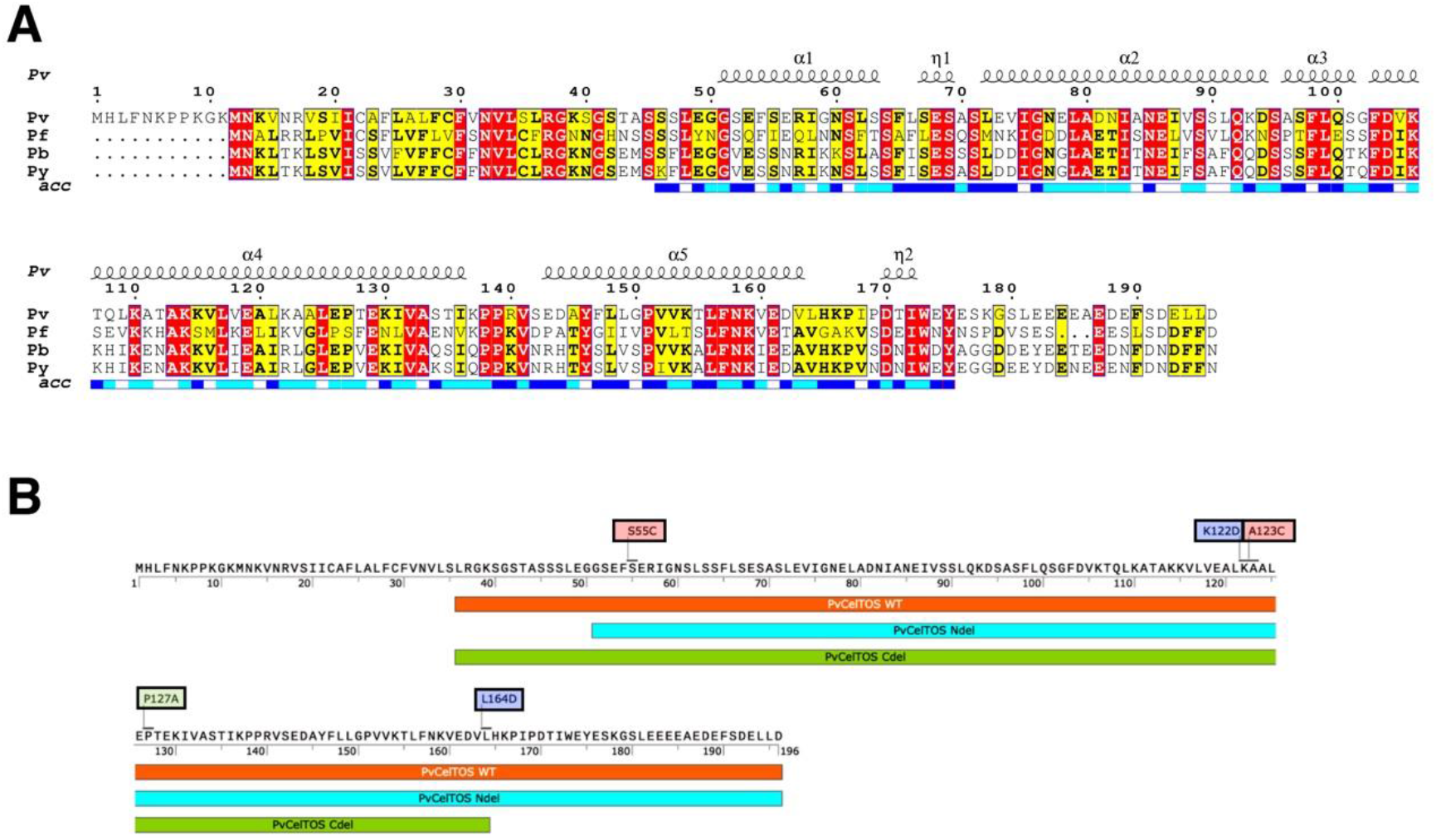
Sequence-based analysis of CelTOS proteins. *(A).* Multiple sequence alignment of CelTOS from different *Plasmodium* species. Pf= *Plasmodium falciparum* 3D7; Pv= *Plasmodium vivax* S01; Pb= *Plasmodium berghei* ANKA; Py= *Plasmodium yoelii yoelli* 17X. Color coding: red highlight– conserved residues; yellow highlight – residues that are conserved among 70% of sequences. Solvent accessibility is mapped below the alignment where blue represents most accessible residues, cyan represents intermittent accessible residues and white represents most accessible residues. The image was generated using Espript server (https://espript.ibcp.fr/ESPript/ESPript/) *(B)* PvCelTOS protein sequence highlighting regions that corresponds to different mutants and were mentioned in the current study. Residues corresponding to single mutants or double mutants are highlighted. P127A mutant = P127A; S-S mutant= S55C and A123C; L164D mutant = L164D and K122D).

**Figure 2 – figure supplement 2:**
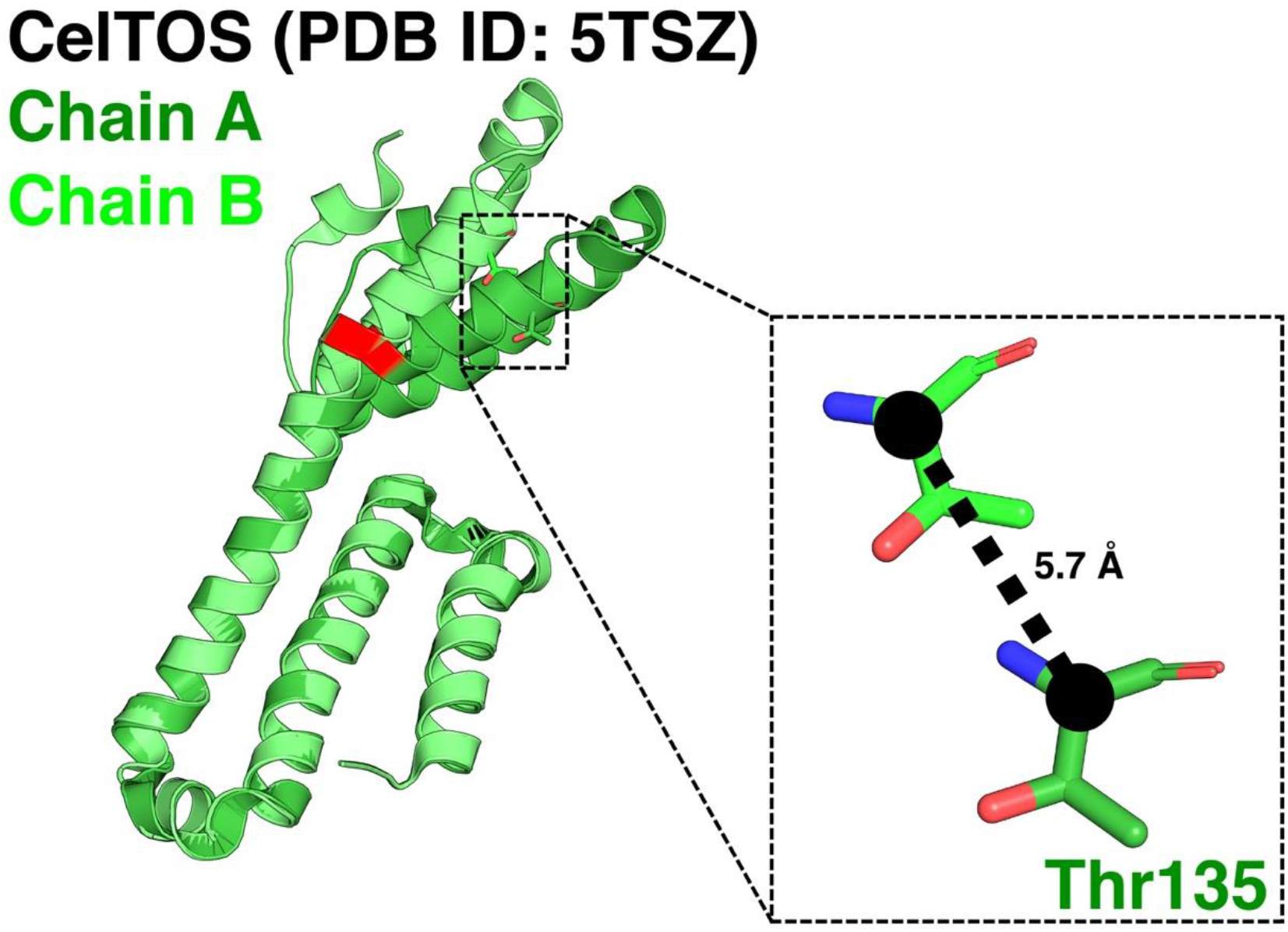
Alignment of the two chains (A, B) of CelTOS structure (PDB ID: 5TSZ). Distance between Cα atoms of Thr135 residues of the two chains is highlighted.

**Figure 2 – figure supplement 3:**
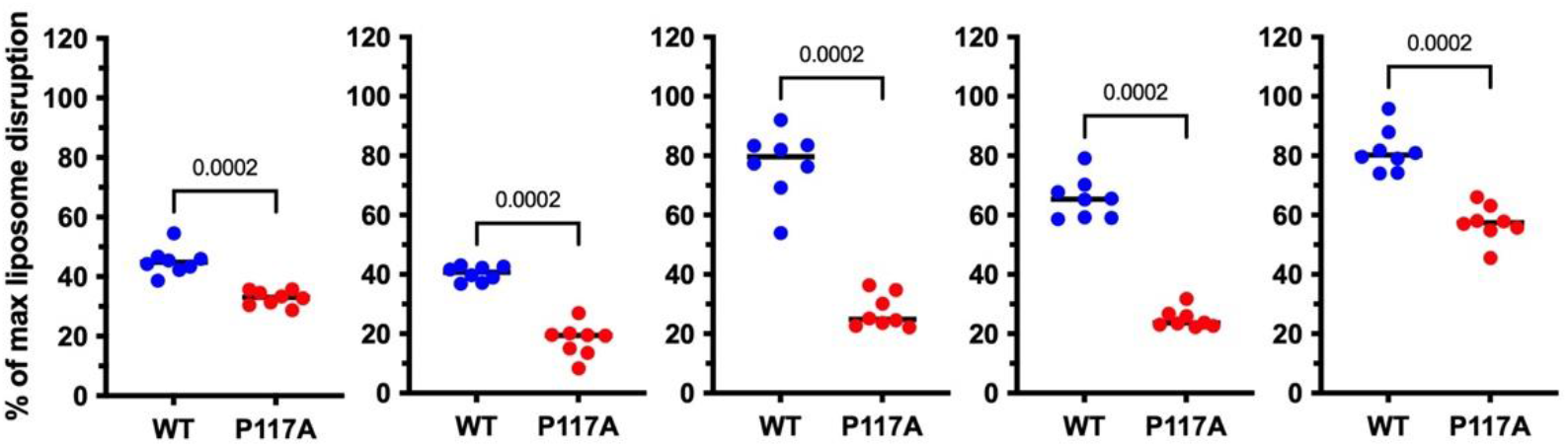
Pore-formation assay of P127A and WT proteins. Each graph represents an individual biological replicate that was performed on a separate day using freshly expressed and purified proteins and freshly prepared liposomes. Lines represent median values. Each biological replicate consists of eight technical replicates. Significance within each biological replicate was determined using an unpaired Mann-Whitney U-test. P-values are shown for each comparison. The y-axis is consistent between each graph and therefore is shown only for the left-most graph. The color-coding is same for each graph and therefore shown only for the right-most graph. The difference in absolute activity is likely due to inherent biological variability derived from each independent batch purification of protein or liposomes. The replicability of the data with independent preparations of proteins and liposomes and performed on different days demonstrate the pore-disruption assay is robust and can distinguish between WT and P117A proteins. The means of each biological replicate were used in Figure 2C to compare WT and P117A proteins across multiple days, batches and preparations.

**Figure 3 – figure supplement 1:**
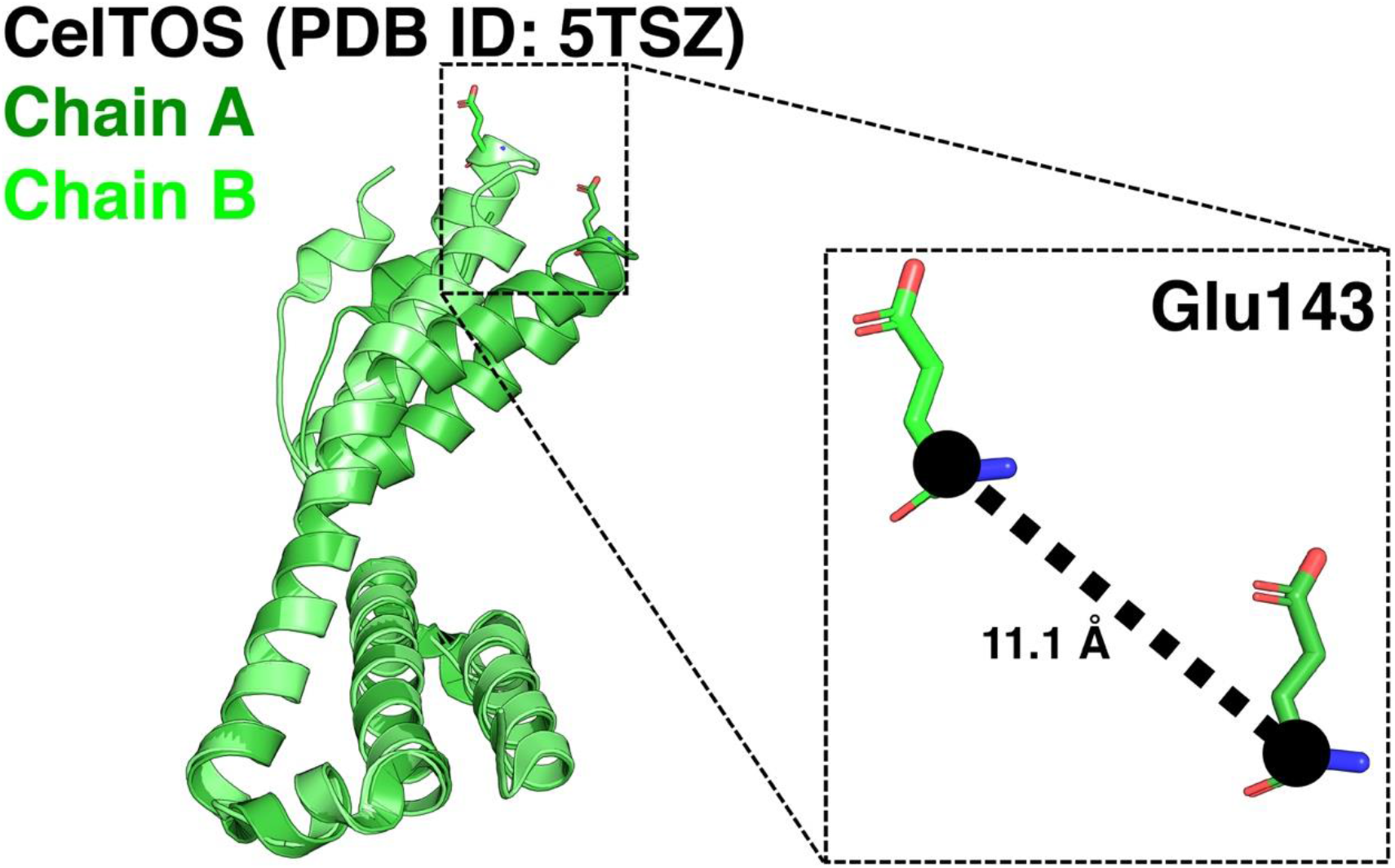
Structural alignment of the two chains (A, B) of CelTOS (PDB ID: 5TSZ). The distance between Cα atoms of Glu143 is shown, representative of flexibility at the C-terminal region.

**Figure 3 – figure supplement 2:**
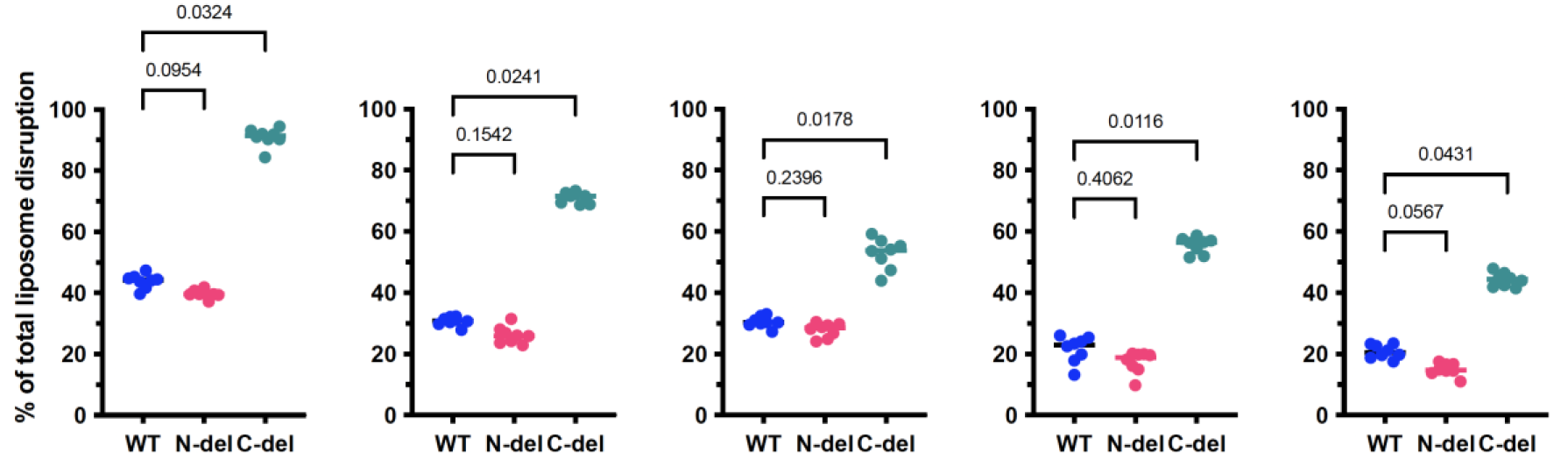
Pore-formation assay of N-del, C-del and WT proteins. Each graph represents an individual biological replicate that was performed on a separate day using freshly expressed and proteins and freshly prepared liposomes. Each biological replicate consists of eight technical replicates. Line represents median values. Significance within each biological replicate was determined using a Kruskal-Wallis analysis and Dunn’s multiple comparison. P-values are shown for each comparison. The y-axis is consistent between each graph and therefore is shown only for the left-most graph. The difference in absolute activity is likely due to inherent biological variability derived from each independent batch purification of protein or liposomes. The replicability of the data with independent preparations of proteins and liposomes, and performed on different days demonstrate the pore-disruption assay is robust and can confidently determine the outcome of different mutations on WT protein. The means of each biological replicate were used in Figure 3B to compare WT, N-del and C-del proteins across multiple days, batches and preparations.

**Figure 4 – figure supplement 1:**
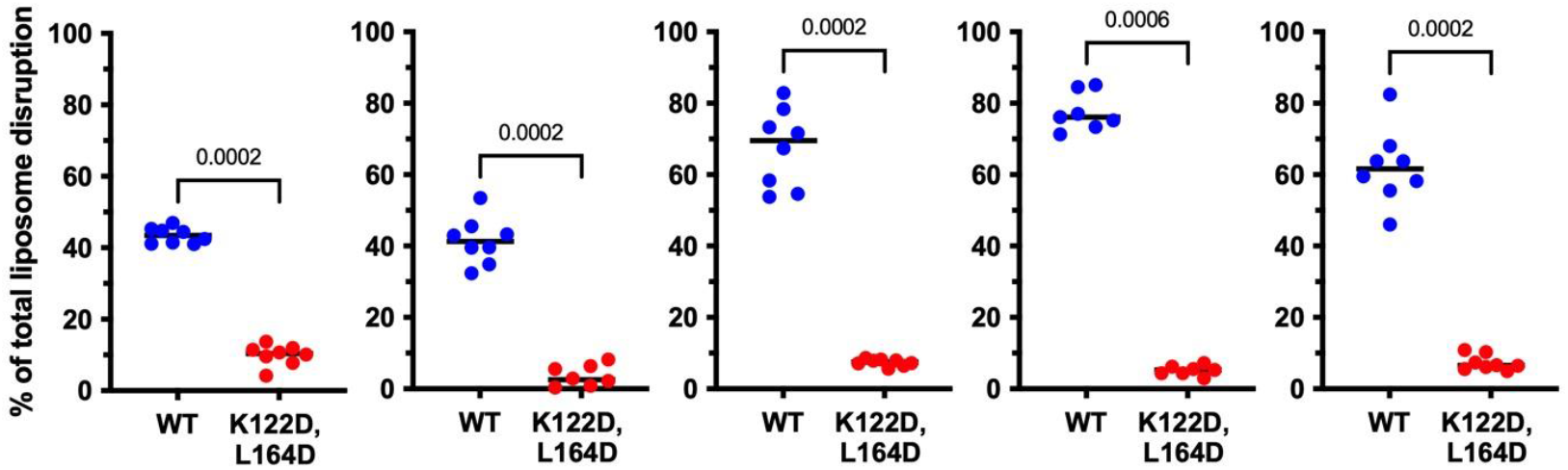
Pore-formation assay of K122D,L164D and WT proteins. Each graph represents an individual biological replicate that was performed on a separate day using freshly expressed and purified proteins and freshly prepared liposomes. Each biological replicate consists of three to four technical replicates. Lines represent median values. Significance within each biological replicate was determined using an unpaired Mann-Whitney U-test. P-values are shown for each comparison. The y-axis is consistent between each graph and therefore is shown only for the left-most graph. The difference in absolute activity is likely due to inherent biological variability derived from each independent batch purification of protein or liposomes. The replicability of the data with independent preparations of proteins and liposomes and performed on different days demonstrate the pore-disruption assay is robust and can distinguish between WT and K122D,L164D mutant. The means of each biological replicate were used in Figure 4E to compare WT and K122D,L164D proteins across multiple days, batches and preparations.

**Figure 4 – figure supplement 2:**
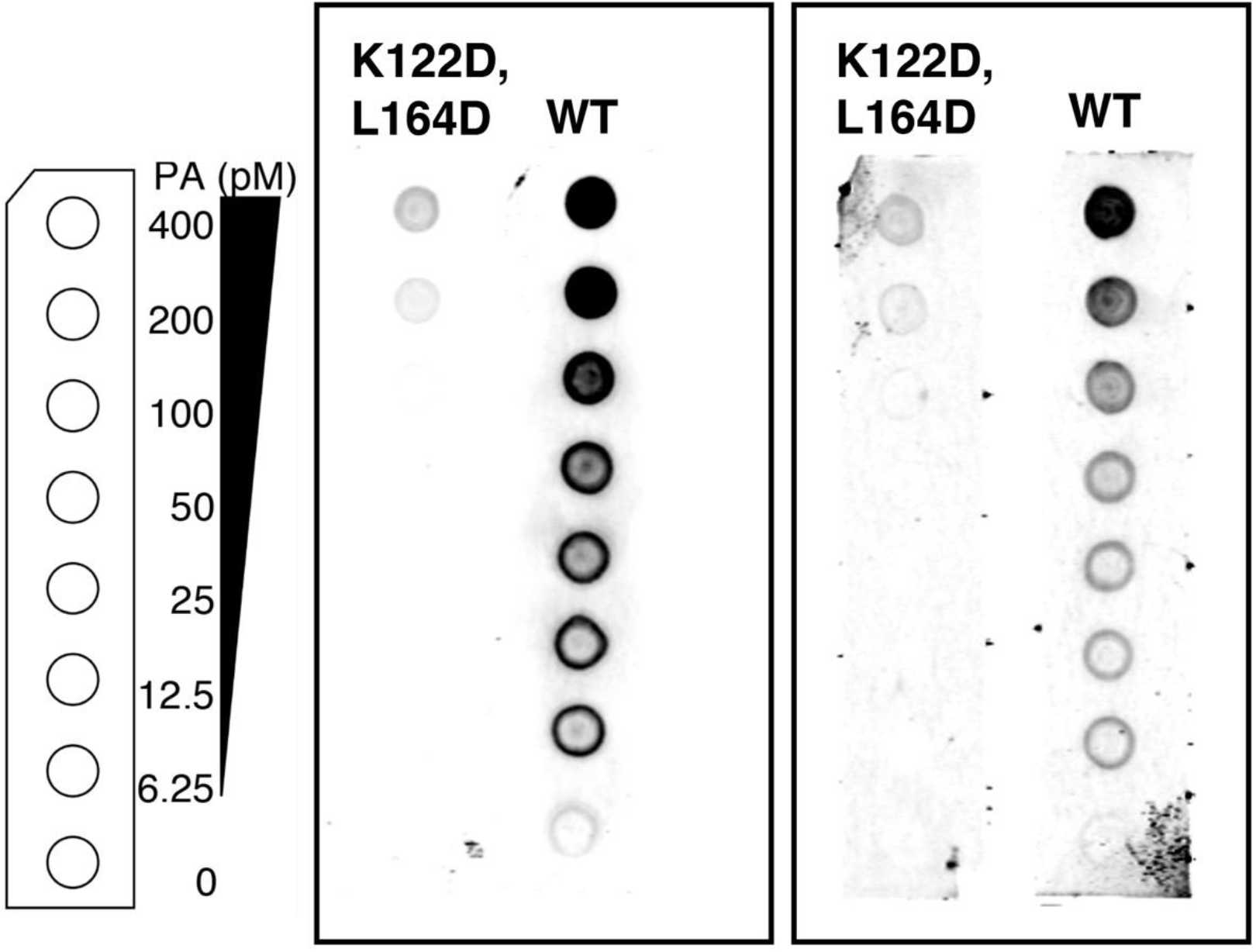
Lipid binding assay of WT and K122D,L164D using phosphatidic acid (PA) strips. Left: a schematic layout of the phosphatidic acid (PA) lipid strips containing a concentration gradient of PA from 400-0 pmol. Right: Two independent biological replicates showing the poor binding interactions between the double mutant and phosphatidic acid (PA).

## References

Anderluh, G., & Lakey, J. H. (2008). Disparate proteins use similar architectures to damage membranes. Trends Biochem Sci, 33(10), 482–490. doi:10.1016/j.tibs.2008.07.004

Bergmann-Leitner, E. S., Hosie, H., Trichilo, J., Deriso, E., Ranallo, R. T., Alefantis, T., … Angov, E. (2013). Self-adjuvanting bacterial vectors expressing pre-erythrocytic antigens induce sterile protection against malaria. Front Immunol, 4, 176. doi:10.3389/fimmu.2013.00176

Bergmann-Leitner, E. S., Legler, P. M., Savranskaya, T., Ockenhouse, C. F., & Angov, E. (2011). Cellular and humoral immune effector mechanisms required for sterile protection against sporozoite challenge induced with the novel malaria vaccine candidate CelTOS. Vaccine, 29(35), 5940–5949. doi:10.1016/j.vaccine.2011.06.053

Bergmann-Leitner, E. S., Mease, R. M., De La Vega, P., Savranskaya, T., Polhemus, M., Ockenhouse, C., & Angov, E. (2010). Immunization with pre-erythrocytic antigen CelTOS from Plasmodium falciparum elicits cross-species protection against heterologous challenge with Plasmodium berghei. PLoS One, 5(8), e12294. doi:10.1371/journal.pone.0012294

Black, R. E., Cousens, S., Johnson, H. L., Lawn, J. E., Rudan, I., Bassani, D. G., … Mathers, C. (2010). Global, regional, and national causes of child mortality in 2008: a systematic analysis. Lancet, 375(9730), 1969–1987. doi:10.1016/s0140-6736(10)60549-1

Case, D. A., Cheatham, T. E., 3rd, Darden, T., Gohlke, H., Luo, R., Merz, K. M., Jr., … Woods, R. J. (2005). The Amber biomolecular simulation programs. J Comput Chem, 26(16), 1668–1688. doi:10.1002/jcc.20290

Cingolani, G., Petosa, C., Weis, K., & Muller, C. W. (1999). Structure of importin-beta bound to the IBB domain of importin-alpha. Nature, 399(6733), 221–229. doi:10.1038/20367

Cosentino, K., & García-Sáez, A. J. (2017). Bax and Bak Pores: Are We Closing the Circle? Trends Cell Biol, 27(4), 266–275. doi:10.1016/j.tcb.2016.11.004

Craig, D. B., & Dombkowski, A. A. (2013). Disulfide by Design 2.0: a web-based tool for disulfide engineering in proteins. BMC Bioinformatics, 14, 346. doi:10.1186/1471-2105-14-346

Dal Peraro, M., & van der Goot, F. G. (2016). Pore-forming toxins: ancient, but never really out of fashion. Nat Rev Microbiol, 14(2), 77–92. doi:10.1038/nrmicro.2015.3

Darden, T., York, D., & Pedersen, L. (1993). Particle mesh Ewald: An N·log(N) method for Ewald sums in large systems. doi:1.464397

Doolan, D. L., Southwood, S., Freilich, D. A., Sidney, J., Graber, N. L., Shatney, L., … Sette, A. (2003). Identification of Plasmodium falciparum antigens by antigenic analysis of genomic and proteomic data. Proc Natl Acad Sci U S A, 100(17), 9952–9957. doi:10.1073/pnas.1633254100

Espinosa, D. A., Vega-Rodriguez, J., Flores-Garcia, Y., Noe, A. R., Muñoz, C., Coleman, R., … Gutierrez, G. M. (2017). The Plasmodium falciparum Cell-Traversal Protein for Ookinetes and Sporozoites as a Candidate for Preerythrocytic and Transmission-Blocking Vaccines. Infect Immun, 85(2). doi:10.1128/IAI.00498-16

Fraser, S. A., Karimi, R., Michalak, M., & Hudig, D. (2000). Perforin lytic activity is controlled by calreticulin. The Journal of Immunology, 164(8), 4150–4155.

Gotz, A. W., Williamson, M. J., Xu, D., Poole, D., Le Grand, S., & Walker, R. C. (2012). Routine Microsecond Molecular Dynamics Simulations with AMBER on GPUs. 1. Generalized Born. J Chem Theory Comput, 8(5), 1542–1555. doi:10.1021/ct200909j

Guerra, A. J., & Carruthers, V. B. (2017). Structural Features of Apicomplexan Pore-Forming Proteins and Their Roles in Parasite Cell Traversal and Egress. Toxins (Basel), 9(9). doi:10.3390/toxins9090265

Gunsteren, W. F. v., & Berendsen, H. J. C. (2006). Algorithms for macromolecular dynamics and constraint dynamics. http://dx.doi.org/10.1080/00268977700102571. doi:Molecular Physics, Vol. 34, No. 5, 1311–1327, November 1977

Homer, M. J., Aguilar-Delfin, I., Telford, S. R., 3rd, Krause, P. J., & Persing, D. H. (2000). Babesiosis. Clin Microbiol Rev, 13(3), 451–469.

Hornak, V., Abel, R., Okur, A., Strockbine, B., Roitberg, A., & Simmerling, C. (2006). Comparison of multiple Amber force fields and development of improved protein backbone parameters. Proteins, 65(3), 712–725. doi:10.1002/prot.21123

Humphrey, W., Dalke, A., & Schulten, K. (1996). VMD: visual molecular dynamics. J Mol Graph, 14(1), 33–38, 27-38.

Jimah, J. R., Salinas, N. D., Sala-Rabanal, M., Jones, N. G., Sibley, L. D., Nichols, C. G.,. Tolia, N. H. (2016). Malaria parasite CelTOS targets the inner leaflet of cell membranes for pore-dependent disruption. Elife, 5. doi:10.7554/eLife.20621

Jimah, J. R., Schlesinger, P. H., & Tolia, N. H. (2017). Liposome Disruption Assay to Examine Lytic Properties of Biomolecules. Bio Protoc, 7(15). doi:10.21769/BioProtoc.2433

Kariu, T., Ishino, T., Yano, K., Chinzei, Y., & Yuda, M. (2006). CelTOS, a novel malarial protein that mediates transmission to mosquito and vertebrate hosts. Mol Microbiol, 59(5), 1369–1379. doi:10.1111/j.1365-2958.2005.05024.x

Kiss, P. T., & Baranyai, A. (2011). Sources of the deficiencies in the popular SPC/E and TIP3P models of water. J Chem Phys, 134(5), 054106. doi:10.1063/1.3548869

Kumar, H., & Tolia, N. H. (2019). Getting in: The structural biology of malaria invasion. PLoS pathogens, 15(9), e1007943.

Kumeta, M., Konishi, H. A., Zhang, W., Sakagami, S., & Yoshimura, S. H. (2018). Prolines in the alpha-helix confer the structural flexibility and functional integrity of importin-beta. J Cell Sci, 131(1). doi:10.1242/jcs.206326

Levine, N. D., Corliss, J. O., Cox, F. E., Deroux, G., Grain, J., Honigberg, B. M., … Wallace, F. G. (1980). A newly revised classification of the protozoa. J Protozool, 27(1), 37–58. doi:10.1111/j.1550-7408.1980.tb04228.x

Mullen, G. E., Giersing, B. K., Ajose-Popoola, O., Davis, H. L., Kothe, C., Zhou, H., … Long, C. A. (2006). Enhancement of functional antibody responses to AMA1-C1/Alhydrogel, a Plasmodium falciparum malaria vaccine, with CpG oligodeoxynucleotide. Vaccine, 24(14), 2497–2505. doi:10.1016/j.vaccine.2005.12.034

Nguyen, V. T., Higuchi, H., & Kamio, Y. (2002). Controlling pore assembly of staphylococcal gamma-haemolysin by low temperature and by disulphide bond formation in double-cysteine LukF mutants. Mol Microbiol, 45(6), 1485–1498.

Ntumngia, F. B., Schloegel, J., McHenry, A. M., Barnes, S. J., George, M. T., Kennedy, S., & Adams, J. H. (2013). Immunogenicity of single versus mixed allele vaccines of Plasmodium vivax Duffy binding protein region II. Vaccine, 31(40), 4382–4388. doi:10.1016/j.vaccine.2013.07.002

Parlati, F., Weber, T., McNew, J. A., Westermann, B., Sollner, T. H., & Rothman, J. E. (1999). Rapid and efficient fusion of phospholipid vesicles by the alpha-helical core of a SNARE complex in the absence of an N-terminal regulatory domain. Proc Natl Acad Sci U S A, 96(22), 12565–12570.

Price, R. N., Tjitra, E., Guerra, C. A., Yeung, S., White, N. J., & Anstey, N. M. (2007). Vivax malaria: neglected and not benign. The American journal of tropical medicine and hygiene, 77(6 Suppl), 79–87.

Rodrigues-da-Silva, R. N., Soares, I. F., Lopez-Camacho, C., Martins da Silva, J. H., Perce-da-Silva, D. S., Teva, A., … Lima-Junior, J. D. (2017). Plasmodium vivax Cell-Traversal Protein for Ookinetes and Sporozoites: Naturally Acquired Humoral Immune Response and B-Cell Epitope Mapping in Brazilian Amazon Inhabitants. Front Immunol, 8, 77. doi:10.3389/fimmu.2017.00077

Roth, A., Maher, S. P., Conway, A. J., Ubalee, R., Chaumeau, V., Andolina, C., … Adams, J. H. (2018). A comprehensive model for assessment of liver stage therapies targeting Plasmodium vivax and Plasmodium falciparum. Nat Commun, 9(1), 1837. doi:10.1038/s41467-018-04221-9

Sastry, G. M., Adzhigirey, M., Day, T., Annabhimoju, R., & Sherman, W. (2013). Protein and ligand preparation: parameters, protocols, and influence on virtual screening enrichments. J Comput Aided Mol Des, 27(3), 221–234. doi:10.1007/s10822-013-9644-8

Schlesinger, P. H., Ferdani, R., Liu, J., Pajewska, J., Pajewski, R., Saito, M., … Gokel, G. W. (2002). SCMTR: a chloride-selective, membrane-anchored peptide channel that exhibits voltage gating. J Am Chem Soc, 124(9), 1848–1849. doi:10.1021/ja016784d

Sharma, V., & Wakode, S. (2016). Pharmacophore generation and atom based 3D-QSAR of quinoline derivatives as selective phosphodiesterase 4B inhibitors. RSC Advances, 6(79), 75805–75819. doi:10.1039/C6RA11210B

Shaw, M. K. (2003). Cell invasion by Theileria sporozoites. Trends in parasitology, 19(1), 2–6.

Sherrill, M. K., & Cohn, L. A. (2015). Cytauxzoonosis: Diagnosis and treatment of an emerging disease. J Feline Med Surg, 17(11), 940–948. doi:10.1177/1098612x15610681

Stote, R. H., & Karplus, M. (1995). Zinc binding in proteins and solution: a simple but accurate nonbonded representation. Proteins, 23(1), 12–31. doi:10.1002/prot.340230104

Tanaka, K., Caaveiro, J. M., Morante, K., González-Mañas, J. M., & Tsumoto, K. (2015). Structural basis for self-assembly of a cytolytic pore lined by protein and lipid. Nat Commun, 6, 6337. doi:10.1038/ncomms7337

Tanaka, K., Caaveiro, J. M., Morante, K., González-Mañas, J. M., & Tsumoto, K. (2015). Structural basis for self-assembly of a cytolytic pore lined by protein and lipid. Nature communications, 6(1), 1–11.

Tuerkova, A., Kabelka, I., Králová, T., Sukeník, L., Pokorná, Š., Hof, M., & Vácha, R. (2020). Effect of helical kink in antimicrobial peptides on membrane pore formation. Elife, 9. doi:10.7554/eLife.47946

Valle, A., Perez-Socas, L. B., Canet, L., Hervis, Y. P., de Armas-Guitart, G., Martins-de-Sa, D., … Pazos, I. F. (2018). Self-homodimerization of an actinoporin by disulfide bridging reveals implications for their structure and pore formation. Sci Rep, 8(1), 6614. doi:10.1038/s41598-018-24688-2

Wang, J., Wolf, R. M., Caldwell, J. W., Kollman, P. A., & Case, D. A. (2004). Development and testing of a general amber force field. J Comput Chem, 25(9), 1157–1174. doi:10.1002/jcc.20035

WHO. (2019). World Malaria Report 2019. Retrieved from Geneva:

